# Rbfox1 is required for myofibril development and maintaining fiber-type specific isoform expression in *Drosophila* muscles

**DOI:** 10.1101/2021.05.09.443278

**Authors:** Elena Nikonova, Ketaki Kamble, Amartya Mukherjee, Christiane Barz, Upendra Nongthomba, Maria L. Spletter

## Abstract

Protein isoform transitions confer distinct properties on muscle fibers and are regulated predominantly by differential transcription and alternative splicing. RNA-binding Fox protein 1 (Rbfox1) can affect both transcript levels and splicing, and is known to control skeletal muscle function. However, the detailed mechanisms by which Rbfox1 contributes to normal muscle development and physiology remain obscure. In this study, we report that Rbfox1 contributes to the generation of adult muscle diversity in *Drosophila*. Rbfox1 is differentially expressed in tubular and fibrillar muscle fiber types. RNAi knockdown of Rbfox1 leads to a loss of flight, climbing and jumping ability, as well as eclosion defects. Myofibers in knockdown muscle are frequently torn, and sarcomeres are hypercontracted. These defects arise from mis-regulation of fiber-type specific gene and splice isoform expression, notably loss of an IFM-specific isoform of Troponin-I that is critical for regulating myosin activity. We find that Rbfox1 influences mRNA transcript levels through 1) direct binding of 3’-UTRs of target transcripts as well as 2) through regulation of myogenic transcription factors, including Mef2, Exd and Salm. Moreover, Rbfox1 modulates splice isoform expression through 1) direct regulation of target splice events in structural genes and 2) regulation of the CELF-family RNA-binding protein Bruno1. Our data indicate that cross-regulatory interactions observed between FOX and CELF family RNA-binding proteins in vertebrates are conserved between their counterparts, Rbfox1 and Bruno1 in flies. Rbfox1 thus affects muscle development by regulation of both fiber-type specific gene and gene isoform expression dynamics of identity genes and structural proteins.

## Introduction

Muscles are an ideal model to understand the strategies involved in the generation of diversity within a tissue, as they are developmentally patterned to be equipped with distinct morphologies and to perform diverse functions (Spletter and Schnorrer, 2014). Muscles develop to accommodate a heterogeneous composition of fiber-types with protein isoform-specific signatures that impart distinctive functionalities to meet diverse physiological demands (Armstrong and Phelps, 1984; Bottinelli, 2001; Bottinelli and Reggiani, 2000; Schiaffino and Reggiani, 2011; Schiaffino et al., 2020). Composite muscle fiber profiles are a result of coordinated regulation of gene expression (Black and Olson, 1998; Firulli and Olson, 1997; Majesky, 2007), co-integrated with protein isoform transitions facilitated by alternative splicing (Guo et al., 2010; Kalsotra and Cooper, 2011; Nikonova et al., 2020; Smith et al., 1989), accompanied by post-translational modifications (Anthony et al., 2002; Michele and Campbell, 2003; Nayak and Amrute-Nayak, 2020; Wells et al., 2003). The underlying molecular changes are initially regulated by the intrinsic developmental program (Firulli and Olson, 1997; Kablar and Rudnicki, 2000), and later modulated by nerve stimulation, physiological demands and patho-physiological conditions (Hughes et al., 1993; Pette and Staron, 2001; Pistoni et al., 2010; Schiaffino et al., 2007). The process of protein isoform expression needs to be tightly regulated to have a functionally relevant outcome (Anthony et al., 2002; Black and Olson, 1998; Firulli and Olson, 1997; Guo et al., 2010; Kalsotra and Cooper, 2011; Majesky, 2007; Smith et al., 1989).

Protein isoform expression is regulated by a diverse array of RNA binding proteins (RBPs). RBPs regulate the process of alternative splicing by binding to *cis*-intronic elements in target RNAs to generate the splicing profile of a given cell type (Fu and Ares, 2014; Kalsotra and Cooper, 2011). RBPs can also regulate translation level by binding to 3’-UTR elements and subsequently associating with translation factors, P-granules or components of the nonsense-mediate decay (NMD) pathway (Hentze et al., 2018; Ho et al., 2021; Kishor et al., 2019). RBPs are thus key mediators of eukaryotic genome information during development and are essential for establishing, refining and maintaining tissue and fiber-type specific properties (Grifone et al., 2020; Lunde et al., 2007; Nikonova et al., 2019; Spletter and Schnorrer, 2014). The salience of this function is illustrated by observations that alternative splicing and protein isoform expression patterns are disrupted in diseases from cardiomyopathy to cancer (Bessa et al., 2020; Picchiarelli and Dupuis, 2020; Ravanidis et al., 2018), and that loss of RBP function leads to severe neuromuscular disorders, such as myotonic dystrophy, amyotrophic lateral sclerosis, and spinal motor atrophy (López-Martínez et al., 2020; Nikonova et al., 2019; Picchiarelli and Dupuis, 2020). It is therefore critically important to understand RBP function in detail.

RNA-binding Fox protein 1 (Rbfox1, the first identified member of the FOX family of RBPs), is an RBP with a single, highly-conserved RNA recognition motif (RRM) domain that binds to 5’-UGCAUG-3’ motifs (Auweter et al., 2006; Jin et al., 2003). Rbfox1 binding to introns causes context-dependent exon retention or skipping, depending on if it binds upstream or downstream of an alternative exon (Fukumura et al., 2007; Nakahata and Kawamoto, 2005), while 3’-UTR binding is reported to modulate mRNA stability (Carreira-Rosario et al., 2016). Rbfox1 may additionally influence transcription networks by binding transcriptional regulators (Shukla et al., 2017; Usha and Shashidhara, 2010; Wei et al., 2016). In vertebrates, Rbfox1 has been shown to either cooperatively or competitively regulate splicing with other RBPs such as SUP-12, ASD-1, MBNL1, NOVA, PTBP, CELF1/2 and PSF (Conboy, 2017; Klinck et al., 2014), as well as to be involved in cross-regulatory interactions with CELF and MBNL family proteins (Gazzara et al., 2017; Sellier et al., 2018). This context-dependent nature of Rbfox1 function, as well as integration with other splicing networks, is not yet fully understood.

Rbfox1 plays an important role in regulating the development of both neurons and muscle (Conboy, 2017). It regulates sensory neuron specification in *Drosophila* (Shukla et al., 2017), and in vertebrates is necessary for proper neuronal migration and axonal growth (Hamada et al., 2016), is induced by stress (Amir-Zilberstein et al., 2012), and modulates the splicing of genes involved axonal depolarization (Gehman et al., 2011; Lee et al., 2009). In vertebrate muscle, Rbfox1 binding sites are enriched around developmentally-regulated, alternatively spliced exons in heart (Kalsotra et al., 2008) and Rbfox1 mediated splicing is involved in the regulation of cardiac failure (Gao et al., 2016). Rbfox1 regulates alternative splicing of structural proteins as well as proteins in the calcium signaling pathway in skeletal muscle, (Pedrotti et al., 2015), and is necessary for the maintenance of skeletal muscle mass (Singh et al., 2018). Rbfox1 mutant mice display myofiber and sarcomeric defects and impaired muscle function (Pedrotti et al., 2015), and Rbfox is downregulated in the mouse model of Facio-scapulo humoral dystrophy (FSHD) (Pistoni et al., 2010). The exact role of Rbfox1 in muscle development and physiology is still not well understood, and is moreover complicated by the presence of other FOX family members in vertebrates, notably Rbfox2 (Begg et al., 2020; Conboy, 2017; Singh et al., 2018), that have similar functions.

Invertebrate models, such as *Drosophila*, are particularly useful to investigate the conserved mechanisms of RBP function in muscle (Nikonova et al., 2019). The *Drosophila* genome has a single copy of the *Rbfox1* (*A2BP1*) gene (Kuroyanagi, 2009), making it easier to study Rbfox1 function without the complexities of redundancy. Muscle structure, as well as the mechanism of acto-myosin contractility, is highly conserved (Dasbiswas et al., 2018; Lemke and Schnorrer, 2017), and studies of alternative splicing regulation and fiber-type specific protein isoform function have proven highly informative (Jagla et al., 2017; Jawkar and Nongthomba, 2020; Plantié et al., 2015). *Drosophila* muscles are of two major types: 1) the fibrillar indirect flight muscles (IFMs) comprised of the dorsal longitudinal (DLMs) and dorso-ventral muscles (DVMs), and 2) the tubular muscles, which constitute the rest of the fly muscles. Fibrillar muscles are physiologically similar to vertebrate cardiac muscles (Peckham et al., 1990; Pringle, 1981; Swank et al., 2006), while tubular muscles resemble those of the vertebrate skeletal muscles (de la Pompa et al., 1989; Nikonova et al., 2020). *Drosophila* muscles also have a uniform fiber-type within a muscle fascicle (Bernstein et al., 1993; Spletter and Schnorrer, 2014), removing the complication of heterogeneous muscle fiber composition found in mammalian muscles.

In the present study, we investigated the role of Rbfox1 in muscle diversity and function using *Drosophila* adult muscles. We show that Rbfox1’s role in muscle development is conserved, as it is necessary for the development of both fibrillar and tubular fiber-types. Impairment of Rbfox1 function in the IFMs causes muscle hypercontraction resulting from the mis-splicing and stoichiometric imbalance of structural proteins. We present evidence that Rbfox1 regulates fiber-type specific isoform expression on multiple levels: 1) regulating transcript levels through direct 3’-UTR binding as well as indirectly through regulation of transcription factors including Extradenticle (Exd), Spalt major (Salm) and Myocyte enhancer factor 2 (Mef2) and 2) regulating isoform expression through direct intronic binding near alternative exons, as well as through regulation of and genetic interaction with the CELF-family splicing factor Bruno1 (Bru1). Notably, Rbfox1, Bruno1 and Salm exhibit level-dependent, cross-regulatory interactions in IFMs. This indicates conservation of an ancient regulatory network between FOX and CELF family proteins in muscle, and moreover suggests a feedback mechanism that integrates RNA-regulation into transcriptional refinement of fiber-type identity.

## Results

### Rbfox1 is differentially expressed between tubular and fibrillar muscles

To evaluate the expression pattern of Rbfox1 in *Drosophila* muscle, we used the protein trap *Rbfox1^CC00511^* (Rbfox1-GFP) fly line (Kelso et al., 2004) to track GFP-tagged Rbfox1 protein expression. We first examined the indirect flight muscles (IFMs), and confirmed earlier data (Usha and Shashidhara, 2010) showing that Rbfox1 is expressed on the wing discs of third instar larvae (L3), in a pattern consistent with the myoblasts (Fig. 1 A). Rbfox1 protein is detectable in IFM nuclei at all stages of adult myofiber development: at 24h APF in IFMs undergoing splitting and myoblast fusion (Fig. 1 B), at 40h APF during sarcomere assembly (Fig. 1 C), at 58h and 72h as sarcomeres undergo maturation (Fig. 1 D, E) and in 2-day old adult IFMs (Fig. 1 F). We also detect continual expression of *Rbfox1* in IFMs at the RNA level based on mRNA-Seq data (Fig. 1 G). Interestingly, we observed a dip in Rbfox1 expression levels around 50h APF in the middle of IFM development on both the protein and mRNA levels.

**Figure 1:**
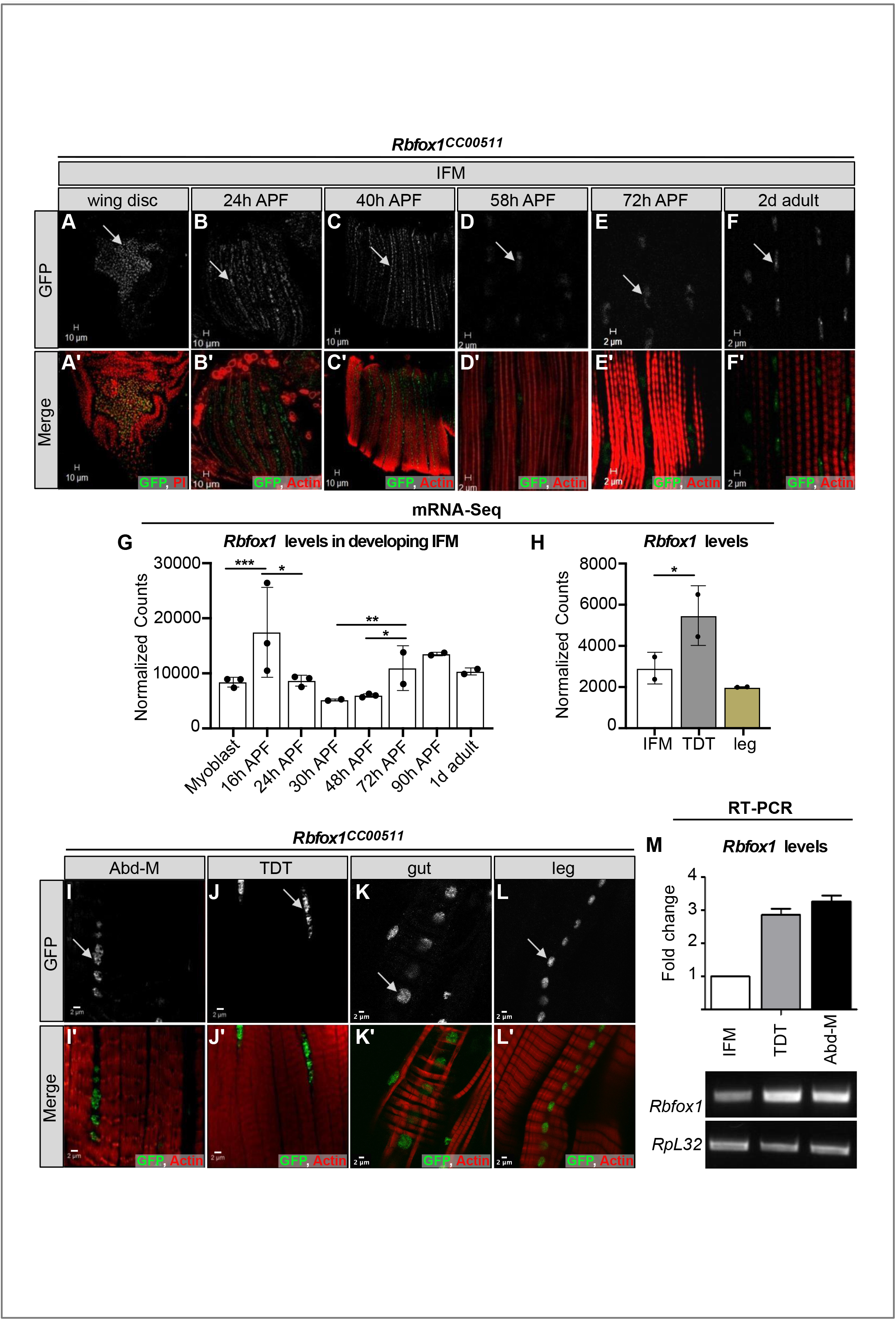
Rbfox1 is differentially expressed between fibrillar and tubular muscle. **A-F.** The *Rbfox1^CC00511^* (Rbfox1-GFP) protein trap line was used to study expression of Rbfox1. **A**) Wing discs of L3 larvae (propidium iodide (PI), red). **B**) IFMs at 24h after puparium formation (APF) show Rbfox1 expression in completely split templates. **C**) IFMs at 40h APF with Rbfox1 expression during initiation of assembly of sarcomere structure. **D** and **E**) IFMs at 58h and 72h APF during sarcomere maturation. **F**) Rbfox1 is expressed in 2-day old adult IFMs. (Arrow indicates GFP positive cells, green; phalloidin stained actin, red; Scale bars = 10 μm.). **G-H**) mRNA-Seq data from *w^1118^* reported as normalized counts show differential expression of Rbfox1 across IFM development (G) and between 1d adult fiber types (H). Significance levels based on DESeq2 analysis (* p < 0.01, ** p < 0.001, *** p< 0.0001). **I-L**) Confocal microscopy of the *Rbfox1*–GFP (*Rbfox1^CC00511^*) line shows Rbfox1 expression in adult tubular muscles including abdominal muscles (Abd-M), TDT, gut and leg (GFP, green; phalloidin stained actin, red). Scale bars = 2 μm. **M**) qPCR and representative semi-quantitative gel images showing relative expression of *Rbfox1* at the mRNA level in adult *Canton-S* across muscle fiber types. *RpL32,* also known as *RP49*, was used as a normalizing control.

We next examined Rbfox1 expression in other types of somatic muscle. Rbfox1-GFP can be detected in the nuclei of all muscles examined, including the tubular abdominal muscles (Abd-M), tergal depressor of the trochanter (TDT or jump muscle), gut and leg muscles (Fig. 1 I-L). Likewise, *Rbfox1* mRNA is detected in all muscles tested, including IFM, TDT, legs and abdomen (Fig. 1 H, M, Fig. S1 A, C). *Rbfox1* mRNA is expressed at significantly higher levels in tubular TDT than in fibrillar IFMs, as revealed by mRNA-Seq (Fig. 1 H) and RT-PCR (Fig. 1 M, Fig. S1 C), and may display preferential exon use between these two fiber types (Fig. S1 B). As leg muscle and Abd-M samples cannot be dissected to the same purity as IFM and TDT, mRNA levels in these samples may not accurately represent muscle-specific *Rbfox1* expression. Taken together, these data demonstrate that although *Rbfox1* is expressed in all types of muscle in *Drosophila,* the expression level is regulated both in a temporal and muscle-type specific manner.

### Rbfox1 function in muscle is necessary for viability and pupal eclosion

To evaluate Rbfox1 function in muscle development, we tested several tools to reduce Rbfox1 levels. We used the deGradFP system, which was developed to specifically target GFP fused proteins (Caussinus et al., 2012), to knockdown *Rbfox1^CC00511^* (Rbfox1-GFP). We also used three RNAi hairpins targeting *Rbfox1*, including *Rbfox1*-RNAi (Usha and Shashidhara, 2010), *Rbfox1-*IR^27286^ and *Rbfox1-*IR*^KK110518^* (Nikonova et al., 2019) (Fig. S1 A). Temporal and spatial regulation of these tools allowed us to evaluate Rbfox1 phenotypes under experimental conditions with different levels of Rbfox1 knockdown.

We started by inducing deGradFP using the constitutive muscle driver Mef2-Gal4, which resulted in early lethality (Fig. 2 A). To restrict knockdown specifically to development of the adult muscles and avoid lethality, we combined Mef2-Gal4 driven Rbfox1^CC00511^-deGradFP and *Rbfox1-*RNAi with *tubulin-Gal80^ts^* and performed a temperature shift from 18 °C to 29 °C at late L3. Temperature-shifted deGradFP flies were pupal lethal and failed to eclose (Fig. 2 A, C). *Rbfox1-*RNAi was less severe, and around 70% of pupae were able to eclose (Fig. 2 A). We confirmed *Rbfox1* knockdown by qPCR (Fig. S1 D). Mef2-Gal4 driven knockdown with *Rbfox1-*IR*^KK110518^* was pupal lethal, and larval lethal when combined with UAS-Dcr2 or driven with Act5c-Gal4 (Fig. 2 A, B). *Rbfox1-*IR*^27286^* was the weakest hairpin, as more than 80% of flies eclosed when crossed to Act5c-Gal4 or Mef2-Gal4. When combined with UAS-Dcr2, most *Rbfox1-*IR^27286^ flies eclosed at 22 °C, but were pupal lethal at 25 °C and 27 °C (Fig. 2 B). We confirmed the level of knockdown by semi-quantitative RT-PCR (Fig. S1 E). We thus are able to tune the expression level of Rbfox1 in muscle and established a knockdown series ordered from strongest to weakest: deGradFP > *Rbfox1-*IR*^KK110518^* > *Rbfox1*-RNAi > *Rbfox1-*IR^27286^. We conclude that Rbfox1 function in muscle is required for viability, as the strongest muscle-specific knockdown conditions resulted in early lethality. Rbfox1 is further required during adult muscle development, as weaker knockdown efficiencies resulted in pupal lethality, notably due to eclosion defects.

**Figure 2:**
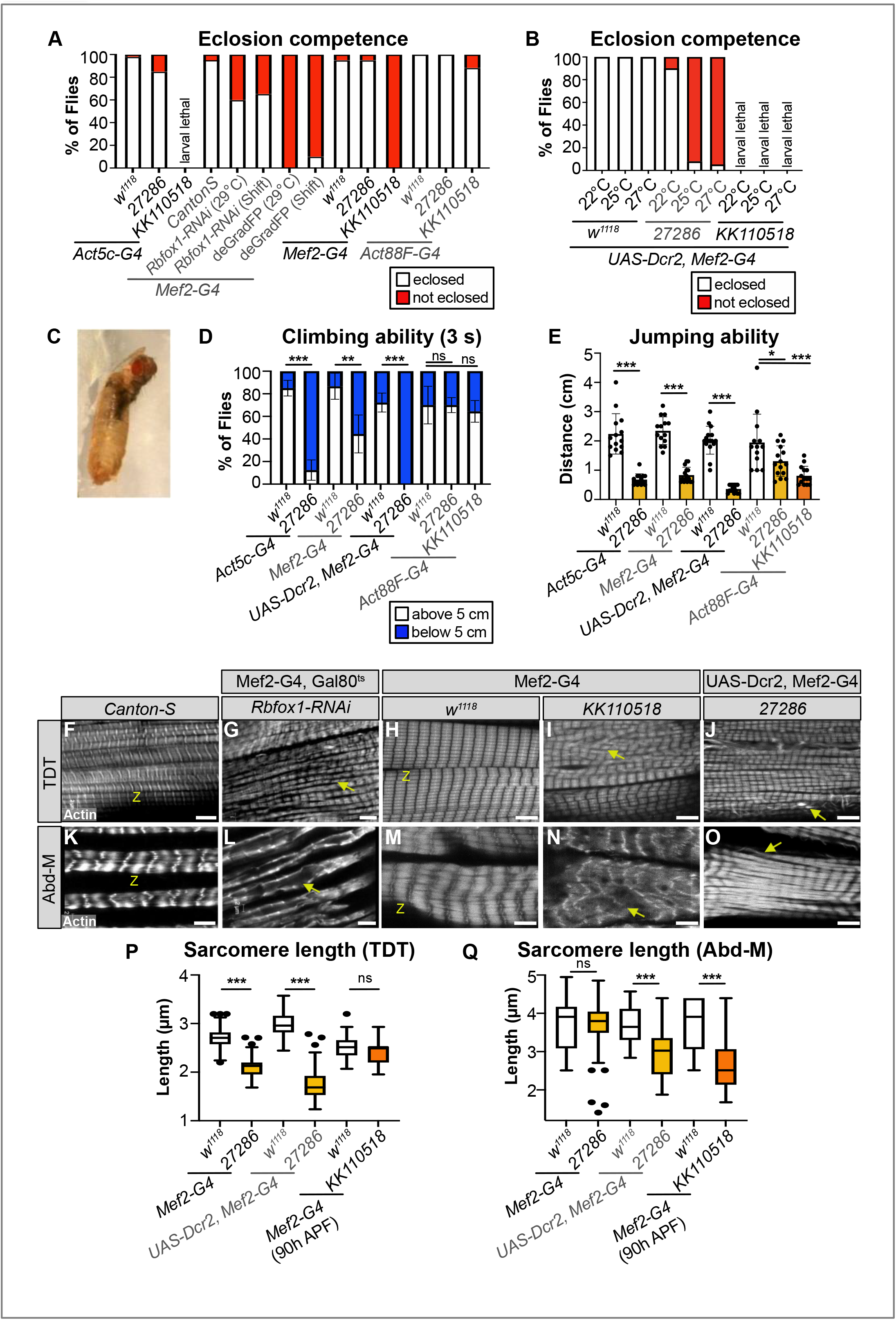
Rbfox1 is necessary for tubular TDT and Abd-M development. **A)** Quantification of the percent of pupae that eclose for controls and Rbfox1 knockdown flies. Genotypes as labelled. **B)** Quantification of the percent of pupae that eclose for UAS-Dcr2, Mef2-Gal4 driven *Rbfox1-IR^27286^*and *Rbfox1-IR^KK110518^* knockdown at 22 °C, 25 °C and 27 °C. **C)** Representative image of the eclosion defect in *Rbfox1-*RNAi. **D)** Quantification of climbing ability measured by how many flies are able to climb 5 centimetres (cm) in 3 seconds (s). **E)** Quantification of jumping ability measured as the distance in cm a startled fly can jump. **F-O)** Single plane confocal images showing myofibril and sarcomere morphology of the TDT (F-J) and Abd-M (K-O). Myofibril structure is altered in *Rbfox1* knockdown conditions, including disorganized myofibril structure (arrow in G, I), frayed myofibrils (arrow in J, O), and loss of sarcomere architecture (arrow in L, N). “Z” indicates z-discs. Scale bars = 5 μm. **P)** Quantification of sarcomere length in TDT. **Q)** Quantification of sarcomere length in Abd-M. Error bars show standard deviation. Significance in D, E, P, Q determined by ANOVA and post-hoc Tukey (not significant, ns; *= p < 0.05; ** = p < 0.01; *** = p < 0.001).

### Rbfox1 contributes to tubular muscle development and function

To determine if Rbfox1 is required in tubular muscles, as suggested by the eclosion defect, we investigated tubular muscle structure and function. We first assayed climbing ability by evaluating how many adult flies were able to climb 5 centimetres (cm) in 3 seconds. We tested *Rbfox1-*IR^27286^ flies driven with Act5c-Gal4 and Mef2-Gal4 at 27 °C, and with UAS-Dcr2, Mef2-Gal4 at 22 °C, as we could obtain surviving adults from these conditions. Flies with reduced Rbfox1 levels were poor climbers (Fig. 2 D), indicating impairment in tubular leg muscle function. We did not observe climbing defects when we performed knockdown with Act88F-Gal4 (Fig. 2 D), which is largely restricted to the fibrillar flight muscles. To assess functional defects in tubular TDT muscle, we then assayed jumping ability by measuring the distance a startled fly can jump. Decreased levels of Rbfox1 severely impaired jumping ability (Fig. 2 E); while control flies on average jumped a distance of around 2 cm, knockdown flies jumped under 1 cm. We also saw defective jumping in Act88F-Gal4 driven Rbfox1 knockdown, and phenotypic severity was dependent on the strength of knockdown (Fig. 2 E). This may reflect a weak expression of the driver in jump muscle, or expression at an earlier point in TDT development. Together, these data indicate that a decrease in Rbfox1 levels results in behaviour defects associated with impaired tubular muscle function.

We next examined tubular muscle structure using confocal microscopy. We observed severe disruptions in sarcomere and myofibril structure in both TDT and Abd-M depending on the strength of Rbfox1 knockdown (Fig. 2 F-O). TDT myofibrils were frayed and severely disorganized after knockdown with all three RNAi hairpins (Fig. 2 F-J). Although nuclei were still organized in the center of the TDT myofibers, the cytoplasmic space between the nuclei was often invaded by myofibrils in knockdown conditions (Fig. S1 H-J). In severe examples, TDT fibers were atrophic and severely degraded (Fig. S1 P). The TDT sarcomeres were significantly shorter in 1d adult flies with Mef2-Gal4 driven *Rbfox1-*IR^27286^ (2.11 ± 0.21 μm versus 2.71 ± 0.19 μm in control, p-value < 0.001) and this was enhanced in the presence of Dcr2 (1.76 ± 0.31 μm versus 2.98 ± 0.26 μm in control, p-value < 0.001). Sarcomeres were not significantly shorter at 90h APF with the stronger Mef2-Gal4 driven *Rbfox1-*IR^KK110518^ (2.43 ± 0.27 μm versus 2.52 ± 0.24 μm in control, p-value = 0.7413) (Fig. 2 P). Classic hypercontraction mutants in IFMs display a similar phenotype, where mis-regulated Myosin activity leads to sarcomere shortening after eclosion (Nongthomba et al., 2003).

We observed similar defects in Abd-M sarcomere and myofibril structure after Rbfox1 knockdown (Fig. 2 K-O). Knockdown with *Rbfox1-*RNAi during adult muscle development led to loss of sarcomere architecture (Fig. 2 L). In *Rbfox1-*IR^27286^ and *Rbfox1-*IR^KK110518^ knockdown animals, Abd-M myofibers were often torn (Fig. 2 M-O) or degraded (Fig. S1 Q). Myofibrils were disorganized, at times invading the center of the fiber (Fig. S1 M-O), and laterally-aligned Z-discs were poorly organized (Fig. 2 M-O). Abd-M sarcomeres in 1d adults with Dcr2, Mef2-Gal4 driven *Rbfox1-*IR^27286^ were significantly shorter than controls (2.99 ± 0.64 μm versus 3.70 ± 0.47 μm in control, p-value < 0.001), and were already significantly shorter at 90h in Mef2-Gal4 driven *Rbfox1-*IR^KK110518^ (2.71 ± 0.83 μm versus 3.74 ± 0.64 μm in control, p-value < 0.001) (Fig. 2 Q). Overall, the observed phenotypes in tubular TDT and Abd-M are consistent between independent RNAi hairpins and phenotypic severity increases with increasing strength of Rbfox1 knockdown. Taken together, the defects in eclosion, climbing, jumping and tubular myofiber morphology indicate a general requirement for Rbfox1 in tubular muscle development.

### Knockdown of *Rbfox1* leads to muscle hypercontraction in the IFMs

We next evaluated Rbfox1 function in fibrillar indirect flight muscle (IFMs). We were able to obtain surviving adults from pupal-restricted *Rbfox1*-RNAi, and noted these flies were completely flightless (Fig. 3 A). In agreement with our previous results (Nikonova et al., 2019), we also found that surviving adults from all *Rbfox1-*IR^27286^ crosses, as well as flies with IFM-restricted, Act88F-Gal4 driven *Rbfox1-*IR*^KK110518^* had impaired flight ability (Fig. 3 B). The strength of the flight defect increased with the strength of the Rbfox1 knockdown and was consistent across all three RNAi hairpins tested.

**Figure 3:**
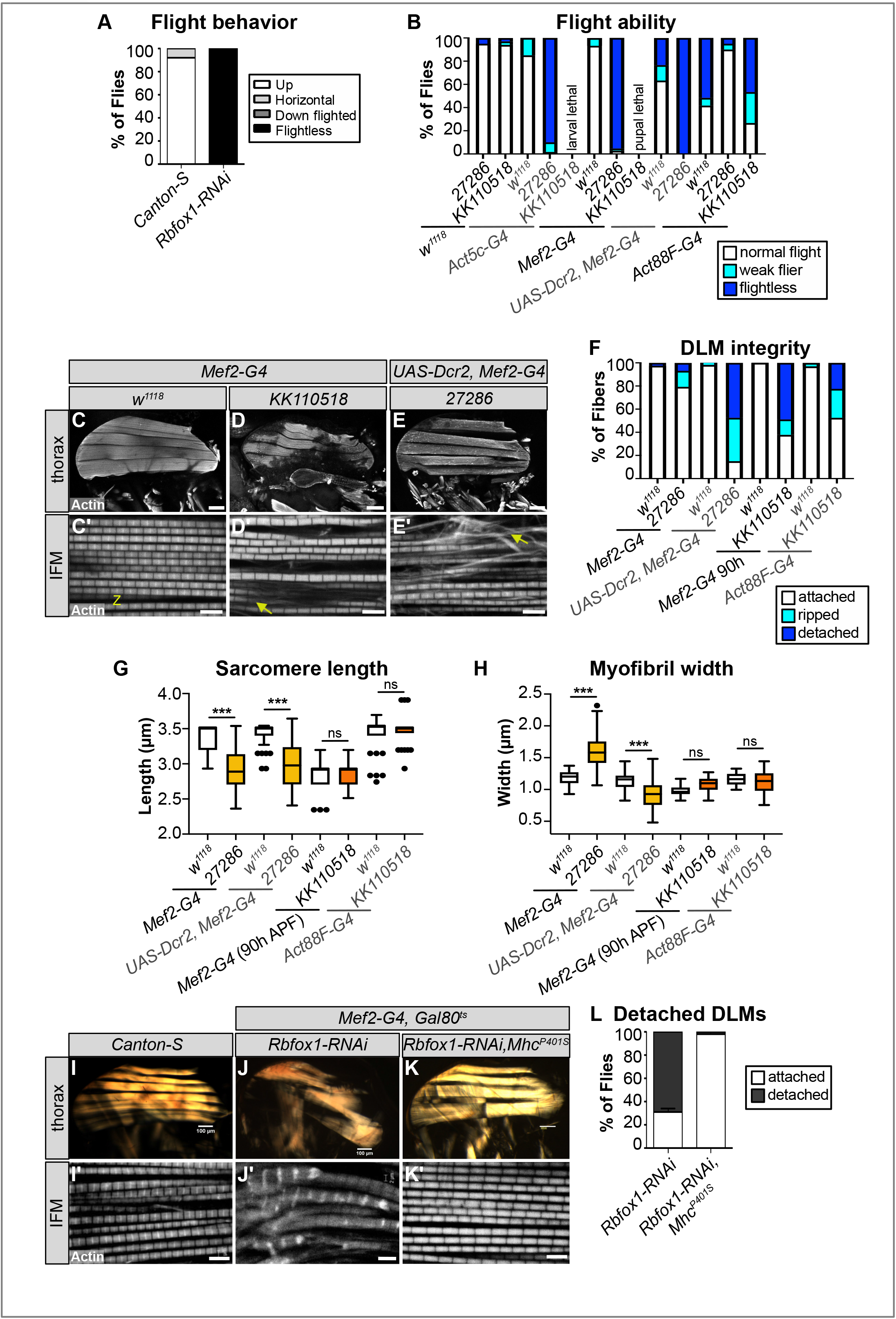
Rbfox1 knockdown results in IFM myofibril defects and hypercontraction-mediated myofiber loss. **A-B)** Quantification of flight ability after Rbfox1 knockdown. Genotypes as noted. **C-E)** Confocal Z-stack images (C-E) of IFM myofiber structure (Scale bars = 5 μm) and single plane images (C’-E’) of myofibril and sarcomere structure after *Rbfox1* knockdown. Note the short sarcomeres and frayed myofibrils (arrow in D’, E’). **F)** Quantification of myofiber ripping and detachment phenotypes in C-E. **G-H)** Quantification of IFM sarcomere length and myofibril width in C’-E’. Error bars show standard deviation. Significance determined by ANOVA and post-hoc Tukey (not significant, ns; *= p < 0.05; ** = p < 0.01; *** = p < 0.001). **I-K)** Polarized microscopy images of hemi-thorax from wild type (I), *Rbfox1-*RNAi (J) and *Rbfox1-*RNAi, *Mhc^P401S^* (K) flies. **I’-K’)** Single-plane confocal images showing phalloidin-stained sarcomeric structure from wild type (I’), *Rbfox1-*RNAi (J’) and *Rbfox1-*RNAi, *Mhc^P401S^* (K’) flies. The *Mhc^P401S^* allele suppresses myofiber loss and sarcomere phenotypes. **L)** Quantification of myofiber detachment in J and K.

To determine if the impaired flight was the result of defective muscle structure or function, we examined IFMs using confocal microscopy. Myofibers in thoraxes of 1d old (1d) adult *Rbfox1-*IR^27286^ flies or 90h APF *Rbfox1-*IR*^KK110518^* flies were frequently torn and detached (Fig. 3 C-F). Myofibrils in remaining intact DLM myofibers were frayed and wavy (Fig. 3 C’-E’). Sarcomere length was significantly shorter in 1d adult flies with both Mef2 > *Rbfox1-*IR^27286^ (2.90 ± 0.24 μm versus 3.34 ± 0.20 μm in control, p-value < 0.001) and with UAS-Dcr2, Mef2-Gal4 enhanced knockdown (2.98 ± 0.33 μm versus 3.43 ± 0.16 μm in control, p-value < 0.001) (Fig. 3 G, Fig. S2 A). Myofibril width in Mef2 > *Rbfox1-*IR^27286^ IFMs was significantly thicker than control (1.58 ± 0.25 μm versus 1.18 ± 0.11 μm in control, p-value < 0.001) (Fig. 3 H, Fig. S2B). Myofibril width was actually thinner with UAS-Dcr2, Mef2-Gal4 enhanced knockdown in 1d adults (0.92 ± 0.22 μm versus 1.14 ± 0.12 μm in control, p-value < 0.001), possibly reflecting the increased severity of myofibril fraying and loss. At 90h APF, sarcomeres of *Rbfox1-*IR^27286^ and *Rbfox1-*IR*^KK110518^* flies were not significantly shorter than the control (Fig. 3 G, Fig. S2 A), but myofibrils tended to be thicker (Fig. S2 B). Myofibrils in Act88F-Gal4 mediated knockdown only showed mild defects (Fig. 3 G, H, Fig. S2 C, D) despite adult flies being flight impaired.

We further confirmed the IFM defects with *Rbfox1-*RNAi and *Rbfox1^CC00511^*-deGradFP. When we assessed DLMs of the few *Rbfox1^CC00511^*-deGradFP escapers, we saw tearing or detachment of muscle fibers (Fig. S2 F-H) and defective patterning of the DLM myofibrils, including actin accumulations and sarcomeric defects (Fig. S2 I, J). We visualized DLM fibers from *Rbfox1*-RNAi adult flies under polarized light and also observed tearing and loss of muscle fibers (Fig. 3 I, J, L). Sarcomere cytoarchitecture was severely disrupted, accompanied by the appearance of actin accumulations at the Z-discs, also known as Zebra bodies (Fig. 3 I’, J’). We were unable to attempt a rescue of these defects because *Rbfox1* over-expression with Mef2-Gal4 was lethal. Over-expression of *Rbfox1* from 40h APF using the IFM-specific UH3-Gal4 (Singh et al., 2014) resulted in an IFM phenotype similar to the knockdown, including torn myofibers (Fig. S2 E) and thin, frayed or torn myofibrils with short sarcomeres (Fig. S2 E’). The consistency in phenotype between all three RNAi hairpins and *Rbfox1^CC00511^*-deGradFP, as well as the increased phenotypic severity with stronger knockdown, indicate that Rbfox1 is required for IFM development. Moreover, the decrease in sarcomere length with a corresponding increase in myofibril width in 1d old adults suggests that loss of Rbfox1 results in a hypercontraction phenotype. Interestingly, both *Rbfox1* knockdown and Rbfox1 over-expression produce similar hypercontraction defects.

Previously, hypercontraction has been characterised as the damage caused by mis-regulated acto-myosin interactions, which can result from many factors including mutations in structural proteins, mechanical stress, stoichiometric imbalance and mis-expression of structural protein isoforms (Firdaus et al., 2015; Nongthomba et al., 2003; Nongthomba et al., 2004; Nongthomba et al., 2007). These mis-regulated acto-myosin interactions can be suppressed by a myosin heavy chain allele (*Mhc^P401S^*) that minimizes the force produced by acto-myosin interactions (Nongthomba et al., 2003). Including the *Mhc^P401S^* allele in the *Rbfox1*-RNAi knockdown background restored the structure of IFM myofibers (Fig. 3 K, L) and sarcomeric cytoarchitecture (Fig. 3K’), confirming that the *Rbfox1* knockdown phenotype indeed resulted from muscle hypercontraction. Complicated genetic recombination prevented us from using the *Mhc^P401S^* allele to additionally confirm the hypercontraction phenotype observed in Rbfox1 over-expression IFMs.

### Bioinformatic identification of muscle genes with Rbfox1 binding motifs

FOX1, the vertebrate homologue of Rbfox1, has previously been shown to regulate splicing and its binding site is over-represented in introns flanking muscle specific exons in vertebrates (Brudno et al., 2001). *Drosophila* Rbfox1 recognizes the same (U)GCAUG motif in RNA (Carreira-Rosario et al., 2016); therefore, we performed a bioinformatic search to identify putative RNA targets of Rbfox1 involved in muscle development. We identified 3,312 genes with intronic Rbfox1 binding motifs, as well as 683 and 1,184 genes with Rbfox1 binding motifs in their 5’-UTR or 3’-UTR regions, respectively (Fig. S3 A, Table S1). The presence of intronic motifs identifies possible alternative splicing targets, while UTR motifs may indicate direct regulation of mRNA stability, trafficking or translation. When classified based on their molecular function gene ontology (GO) annotation, many of these genes have binding or catalytic activity, notably including DNA, RNA and actin-binding, and a portion are structural molecules (Fig. S3 B). When we look in previously annotated gene lists (Spletter et al., 2018), around 20% of all RNA-binding proteins, 40% of transcription factors and 60% of sarcomere proteins contain Rbfox1 binding motifs in their introns or UTR regions (Fig. S3 C). Overall, about 30% of genes that have reported RNAi phenotypes in muscle (Schnorrer et al., 2010) and nearly 25% of genes regulated in a fibrillar-specific manner (Spletter et al., 2015) also contain canonical Rbfox1 binding motifs. These genes influencing muscle development are enriched for Biological Process GO terms such as “regulation of transcription, DNA-templated”, “regulation of RNA metabolic process”, “actin cytoskeleton organization” and “sarcomere organization” (Fig. S3 D). We also see enrichment for terms like “signal transduction”, “synapse organization” and “axon guidance,” suggesting that characterized roles for Rbfox1 in neuronal development (Gehman et al., 2011) may also affect the neuro-muscular junction. This strongly suggests that genes important for muscle development are likely targets of Rbfox1 regulation.

We then selected candidate Rbfox1 target genes to verify based on their direct or indirect involvement in muscle contraction, which could explain the sarcomere defects and mis-regulated acto-myosin interactions in the *Rbfox1* knockdown condition. We found that the characterized myogenic transcription factors *extradenticle* (*exd*) and *Myocyte enhancer factor 2* (*Mef2*) contain 3 and 7 Rbfox1 motifs, respectively, both in introns and 3’-UTR regions (Fig. S3 F, G). The RNA-binding protein Bruno1 (Bru1, also called *arrest*), which has previously been shown to regulate fibrillar-specific alternative splicing (Oas et al., 2014; Spletter et al., 2015), contains 42 intronic and 2 5’-UTR Rbfox1 binding motifs (Fig. S3 H). We also noted that putative Rbfox1 targets include proteins with structural molecule activity such as Troponin-I (TnI), which is encoded by the gene *wings up A* (*wupA*). TnI is the inhibitory subunit of the Troponin complex and has an Rbfox1 binding site downstream of exon 6b1 and another in the 3’-UTR (Fig. S3 E). The TnI isoform containing exon 6b1 and exon 3 is reported to be specific to the IFMs and its loss was previously shown to result in hypercontraction (Barbas et al., 1993; Nongthomba et al., 2004). We next proceeded to experimentally validate these candidate genes.

### Rbfox1 regulates expression of structural proteins

To confirm if Rbfox1 regulates target structural proteins including TnI, we checked the expression of TnI in *Rbfox1*-RNAi IFMs. TnI protein levels were significantly upregulated in the IFMs with *Rbfox1* knockdown as assayed by Western Blot (Fig. 4 A, B). Although not significant, *wupA* mRNA levels tend towards upregulation in *Rbfox1*-RNAi IFMs and TDT as assayed by RT-qPCR (Fig. S4 A). Overexpression of Rbfox1 significantly reduced the levels of TnI protein detected by Western Blot in IFMs (Fig. 4 D, E). *wupA* mRNA levels were not significantly changed with Rbfox1 overexpression, but tended towards upregulation (Fig. S4 A). As a control which lacks the Rbfox1 binding site (Table S1), we assessed expression of the flight muscle specific actin (Act88F). Although Act88F levels in IFMs tended toward upregulation, we did not observe statistically significant changes in either Act88F protein (Fig. 4A, C) or mRNA (Fig. S4 B, C, D) levels. Surprisingly, overexpression of Rbfox1 significantly decreased the expression level of Act88F protein (Fig. 4 D, F) and mRNA (Fig. S4 B). In addition, Rbfox1 knockdown in TDT resulted in significantly decreased levels of Act88F mRNA (Fig. S4 C, D). Thus, Rbfox1 negatively regulates expression of structural proteins TnI and Act88F in IFMs, and positively regulates Act88F mRNA levels in TDT.

**Figure 4:**
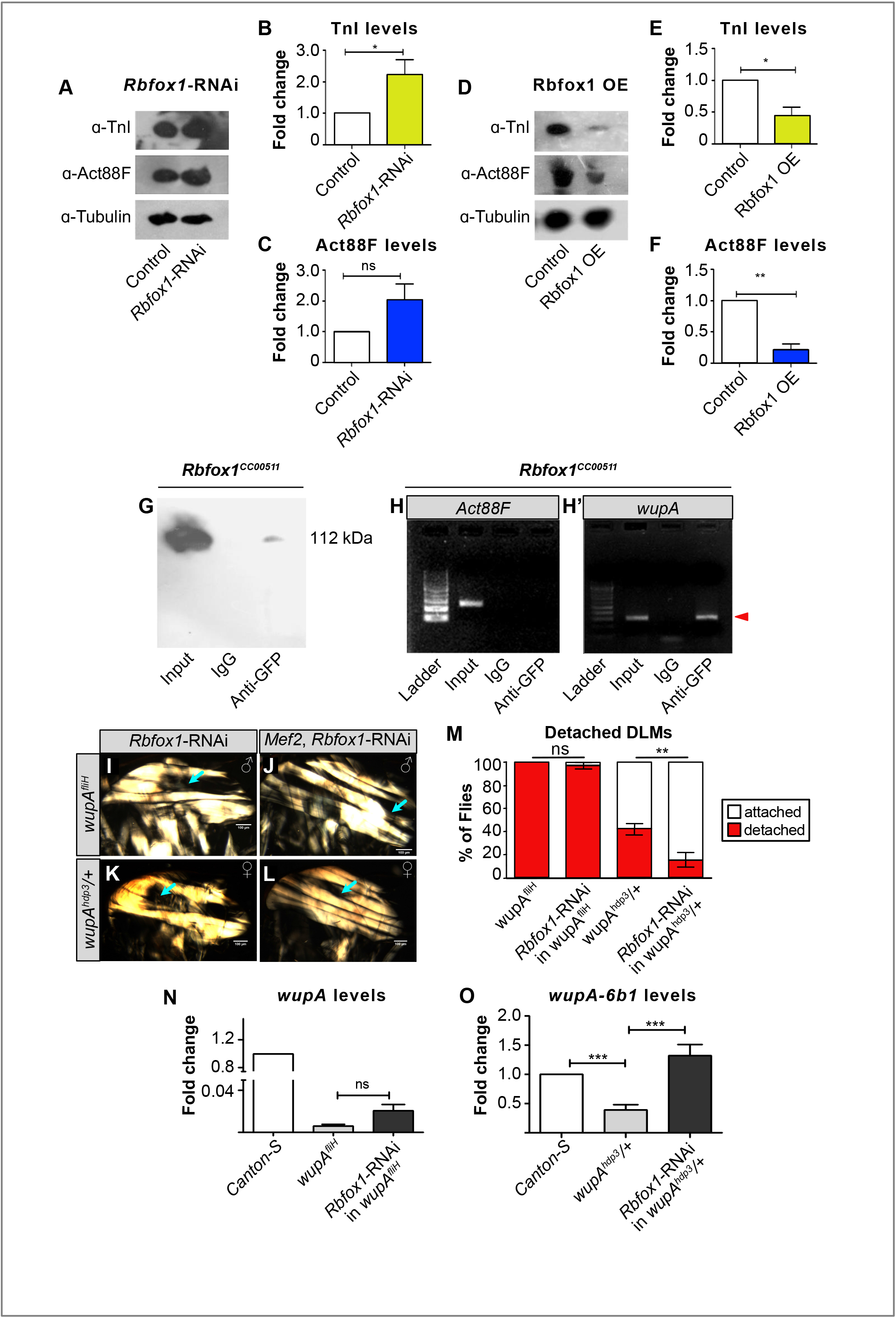
Expression of structural proteins in IFM is regulated by Rbfox1. **A)** Western blot for TnI, Act88F and Tubulin protein levels in *Rbfox1*-RNAi IFM. **B-C)** Quantification of TnI (B) and Act88F (C) expression levels from (A), normalized against Tubulin signal. **D)** Western blot for TnI, Act88F and Tubulin protein levels in IFM with UH3-Gal4 driven Rbfox1 overexpression (Rbfox1 OE). **E-F)** Quantification of TnI (E) and Act88F (F) expression levels from (B), normalized against Tubulin signal. Error bars in B, C, E, F show standard deviation; data from 3 biological replicates. Significance is from paired t-test (not significant, ns; *= p < 0.05; ** = p < 0.01). **G)** Western blot confirming Rbfox1-GFP (*Rbfox1^CC00511^*) is selectively immunoprecipitated with anti-GFP antibody. **H, H’**-Gels showing RNA immunoprecipitation (RIP) followed by RT-PCR from *Rbfox1*-GFP thoraces. mRNA from *Act88F,* which does not have an Rbfox1 binding site, is not detected via RIP (H), while *wupA* (TnI) mRNA can be detected via RIP (red arrowhead, H’), indicating direct Rbfox1 binding. **I-L)** Polarized microscopy images of hemi-thoraxes from *wupA^fliH^* hemizygous males (I), *wupA^fliH^, Rbfox1*-RNAi males (J), *wupA^hdp-3/+^* heterozygous females (K), and *wupA^hdp-3/+^, Rbfox1*-RNAi females (L) with detached IFM myofibers (cyan arrow). Scale bars = 100 μm **M)** Quantification of myofiber attachment in I-L reveals a partial rescue in *wupA^hdp-3/+^, Rbfox1*-RNAi females. Significance is from paired t-test, ** = p < 0.01. **N)** RT-qPCR for *wupA* mRNA transcript levels in IFM from *Canton-S*, *wupA^fliH^*, and *wupA^fliH^, Rbfox1*-RNAi males. **O)** RT-qPCR for *wupA-6b1* mRNA transcript levels in IFM from *Canton-S*, *wupA^hdp-3/+^*, and *wupA^hdp-3/+^, Rbfox1*-RNAi females. Significance is from paired t-test (not significant, ns; *** = p < 0.001).

To determine whether Rbfox1 directly binds *wupA* (TnI) and *Act88F* mRNAs, we performed RNA immunoprecipitation (RIP) to pull down target RNAs bound to Rbfox1. We used the *Rbfox1^CC00511^* (Rbfox1-GFP) fly line and confirmed via Western blot that anti-GFP antibodies could selectively immunoprecipitate Rbfox1-GFP (Fig. 4 G). We then amplified RNA bound to Rbfox1 by RT-PCR with gene-specific primers (Table S2). *Act88F*, which lacks Rbfox1 binding sites and thus served as the negative control, could not be detected after RIP (Fig. 4 H). By contrast*, wupA* (TnI) mRNA was enriched in the RIP with anti-GFP antibodies, but not in the IgG isotype control (Fig. 4 H’). This likely reflects Rbfox1 binding to the motif in the 3’-UTR of *wupA*, as the PCR primers amplify the C-terminal region of the fully-spliced mRNA transcript. Thus, Rbfox1 binds directly to *wupA* mRNA to regulate its expression.

While Rbfox1 binding sites in introns are typically associated with regulation of alternative splicing (Conboy, 2017), a recent study showed that Rbfox1 binds to the UTR region of *Pumilio* mRNA and represses its translation in *Drosophila* ovaries (Carreira-Rosario et al., 2016). To check whether Rbfox1 regulates some target mRNAs such as *wupA* (TnI) post-transcriptionally in IFMs, we looked for interacting partners of Rbfox1. We performed co-immunoprecipitation from *Rbfox1^CC00511^* (Rbfox1-GFP) thoraxes followed by mass spectrometry to identify protein interactors (Fig. S4 E-G). We found that Rbfox1 interacted with the cellular translation machinery including the eukaryotic translation initiation factor eIF4-A and Rent-1 (a regulator of nonsense mediated decay) (Fig. S4 G). These findings suggest that Rbfox1 may regulate translation or direct target mRNAs to nonsense mediated decay through 3’-UTR binding.

### Mis-regulation of TnI contributes to hypercontraction in Rbfox1 knockdown IFMs

We wondered if the hypercontraction phenotype observed after Rbfox1 knockdown and overexpression could be caused by mis-regulation of TnI, thus we performed genetic interaction studies with TnI alleles. It was previously reported that mutations in the TnI encoding *wupA* gene cause hypercontraction in the IFMs. A mutation in the splice site preceding exon 6b1 leads to an IFM-specific null mutant *wupA^hdp-3^* (Barbas et al., 1993), which shows a hypercontraction phenotype in the heterozygous condition (Nongthomba et al., 2004). Another mutant *wupA^fliH^* has a mutation in the Mef2 binding site located in an upstream response element (URE) and results in hypercontracted IFMs with reduced levels of TnI (Firdaus et al., 2015). Since *Rbfox1* knockdown increases TnI levels (Fig. 4 A, B), we knocked down *Rbfox1* in each of the *wupA^fliH^* and *wupA^hdp-3^* mutant backgrounds to see if TnI levels were restored and hypercontraction was rescued. As *wupA^fliH^* is a recessive mutation, hemizygous males were used. *Rbfox1*-RNAi in the *wupA^fliH^* background did not rescue muscle hypercontraction (Fig. 4 I, J, M), and *TnI* levels did not change significantly (Fig. 4 N). However, *Rbfox1*-RNAi in *wupA^hdp-3^* heterozygous mutant female flies showed partially rescued the IFM hypercontraction phenotype and significantly reduced myofiber loss (Fig. 4 K, L, M). We confirmed that *wupA^fliH^* heterozygous mutants have 36-40% of *wupA* mRNA expression compared to *Canton-S* controls, as reported previously (Nongthomba et al., 2004). We also observed that mRNA levels of specifically the *wupA-6b1* transcript were restored when *Rbfox1* was knocked down in *wupA^hdp-3^* mutants (Fig. 4 O). Thus, *Rbfox1* knockdown rescued hypercontraction in *wupA^hdp-3^* but not in *wupA^fliH^* mutants, suggesting that in addition to direct 3’-UTR binding, Rbfox1 may influence muscle-specific splicing of *wupA* (TnI). These results demonstrate that Rbfox1 regulation of TnI expression contributes to the muscle hypercontraction phenotype.

### Rbfox1 regulates splicing factor Bruno1 levels across muscle fiber-types

The mammalian Rbfox1 ortholog FOX1 not only performs tissue specific splicing of target mRNAs during muscle development (Nakahata and Kawamoto, 2005), but is also subject to complex, cross-regulatory interactions with CELF family RNA-binding proteins (Nikonova et al., 2019). One of the top hits in our bioinformatic analysis with 44 Rbfox1 binding motifs was Bruno1 (Bru1) (Fig. 5 A, Fig. S3 H), a CELF1/2 homologue in *Drosophila*. Bru1 has previously been shown to be necessary and sufficient for IFM-specific alternative splicing of structural protein genes including *wupA* (TnI) (Oas et al., 2014; Spletter et al., 2015). This led us to investigate if Rbfox1 might regulate Bru1 in *Drosophila*.

**Figure 5:**
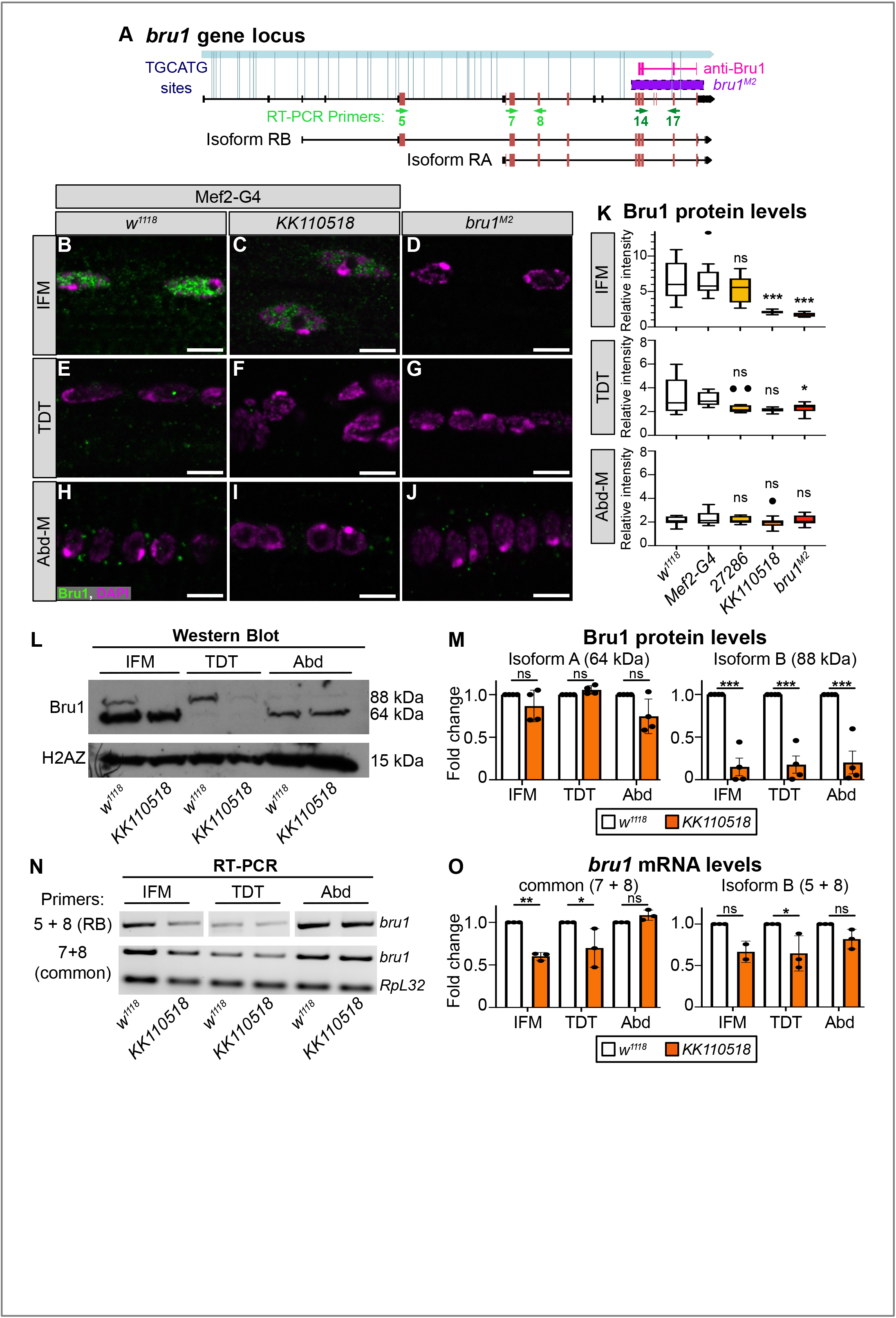
Rbfox1 regulates expression of the RNA-binding protein Bru1. **A)** Diagram of the *bruno1* (*bru1*) locus. The *bru1-RA* and *bru1-RB* isoforms, target region of the rabbit anti-Bru1 antibody (magenta), region deleted in the *bru1^M2^* allele (purple), and TGCATG Rbfox1 binding motifs (light blue) are indicated. Exons, red; UTR, black; RT-PCR primers, green. Not drawn to scale. **B-J)** Confocal images of immunostaining with rabbit anti-Bru1 in IFM (B-D), TDT (E-G) and abdominal muscle (Abd-M) (H-J). Bru1 signal is reduced with *Rbfox1-IR^KK110518^* (C, F, I) and absent in *bru1^M2^* mutant muscle (D, G, J). Bru1, green; DAPI, magenta; Scale bars = 5 μm. **K)** Quantification of Bru1 fluorescence levels in B-J. Significance determined by ANOVA and post-hoc Tukey in comparison to both wild-type (*w^1118^*) and Gal4 alone (*Mef2-Gal4 x w^1118^)* controls (not significant, ns; *= p < 0.05; *** = p < 0.001). **L)** Western blot of Bru1 protein levels in IFM, TDT and abdominal carcass (Abd). Levels of Bru1-PA isoform (at 64 kDa) do not change, while levels of the Bru1-PB isoform (at 88 kDa) decrease in *Rbfox1-IR^KK110518^* muscle. H2AZ was used as a loading control. **M)** Quantification of fold change in band intensity in L, normalized to H2AZ and control IFM expression levels. *w^1118^*, white; *Rbfox1-IR^KK110518^*, red. **N)** Semi-quantitative RT-PCR with primers specific to *bru1-RB* (primers 5 + 8) or common to all *bru1* isoforms (primers 7 + 8). *RpL32* (*RP49*) was used as a control. **O)** Quantification of fold change in band intensity in N, normalized to *RpL32* and control IFM expression levels. Significance in M and O determined by ANOVA and post-hoc Tukey (not significant, ns; *= p < 0.05; ** = p < 0.01, *** = p < 0.001).

We first evaluated Bru1 expression in Rbfox1 knockdown muscle at the protein level using a rabbit polyclonal antibody generated against the divergent domain between RRM2 and RRM3 that should recognize all Bru1 isoforms (Fig. 5 A). In immunostainings of wildtype (*w^1118^*) adult IFMs, Bru1 is strongly expressed and localized to the nucleus (Fig. 5 B). In IFMs from 1d old adult flies with Mef2-Gal4 driven *Rbfox1-*IR^27286^ or in 90 h APF *Rbfox1-*IR*^KK110518^* IFMs, we observed reduced Bru1 staining (Fig. 5 C, K). Bru1 staining is absent in a CRISPR mutant *bru1^M2^* that removes the divergent domain, demonstrating antibody specificity (Fig. 5 A, D, G, J, K). We were able to detect extremely low levels of mostly cytoplasmic Bru1 in wildtype TDT, and this staining was lost after Rbfox1 knockdown and undetectable in *bru1^M2^* mutant TDT (Fig. 5 E-G, K). There was no Bru1 signal detectable above background in any Abd-M samples (Fig. 5 H-J, K). Thus, Rbfox1 knockdown leads to a reduction of Bru1 protein levels in IFMs and TDT.

We next assessed Bru1 protein levels using Western blot. In IFMs dissected from wildtype flies, we consistently observed a band at 64 kDa and a second at 88 kDa (Fig. 5 L), presumably corresponding to the Bru1-PA and Bru1-PB protein isoforms produced from the *bru1-RA* and *bru1-RB* mRNA transcripts, respectively (Fig. 5 A). TDT predominantly expresses Bru1-PB, while weak expression of Bru1-PA is detected in Abd (Fig. 5 L). Bru1-PB was significantly reduced in IFMs and TDT from *Rbfox1-*IR*^KK110518^* flies, while the Bru1-PA isoform was largely unaffected (Fig. 5 L, M). At the mRNA level, semi-quantitative RT-PCR using primers targeting a region common to both isoforms revealed a decrease in *bru1* levels in both IFMs and TDT (Fig. 5 N, O). Levels of specifically the *bru1-RB* transcript were also reduced in both IFMs and TDT (Fig. 5 N, O). We performed RIP to determine if Rbfox1 regulation of *bru1* mRNA is direct and indeed could detect *bru1* RNA bound to Rbfox1-GFP, but we are unable to resolve the specific transcript or distinguish between mature mRNA or partially spliced pre-mRNA in the bound fraction (Fig. S4 H). We also tested a GFP reporter built with the promoter region upstream of *bru1-RA*, but observed no change in GFP expression in *Rbfox1-*RNAi IFMs (Fig. S4 I-J). This is either due to differential regulation of the *bru1-RA* and *bru1-RB* isoforms, or more likely due to regulation of *bru1* RNA processing rather than transcription. We conclude that Rbfox1 regulates levels of Bru1 in both fibrillar IFMs and tubular TDT, and preferentially targets the *bru1-RB* isoform.

### Rbfox1 and Bru1 cross-regulatory interactions are expression level dependent

In vertebrates, members of the FOX and CELF families of RNA-binding proteins display complex cross-regulatory interactions, and we were curious if these interactions are evolutionarily conserved in flies. First, Rbfox1 has been shown to auto-regulate its own expression (Damianov and Black, 2010), and indeed we find 35 Rbfox1 binding motifs in Rbfox1 introns (Figure S5 A). Although strong knockdown with *Rbfox1-*IR*^KK110518^* and Dcr2 enhanced *Rbfox1-*IR^27286^ significantly decreases levels of *Rbfox1* mRNA, a weaker knockdown with *Rbfox1-*IR^27286^ tends towards increased *Rbfox1* levels (Fig. S1 E), suggesting de-repression of a negative feedback loop. Second, our data with a strong *Rbfox1-*IR*^KK110518^* knockdown indicate that Rbfox1 can positively regulate Bru1 protein levels (Fig. 5). However, in mRNA-Seq data from Spletter et al., (2018), we observed that *Rbfox1* and *bru1* have opposite temporal mRNA expression profiles across IFM development, suggesting that Bru1 levels are high when Rbfox1 levels are low (Fig. S5 B). To evaluate if Rbfox1 expression levels might alter the valence of the regulatory interaction with Bru1, we took advantage of our RNAi knockdown series. Indeed, weaker knockdown conditions with *Rbfox1-*IR^27286^ as well as *Rbfox1-*RNAi resulted in increased levels of *bru1* mRNA as well as the *bru1-RB* transcript in IFMs (Fig. S5 C, D). Correspondingly, we see a trend towards increased protein-level expression of Bru1-PA in *Rbfox1-*IR^27286^ IFMs, although levels of both Bru1-PA and Bru1-PB do not change significantly (Fig. S5 E, F). Mef2-Gal4 driven Rbfox1 overexpression, accomplished by a temperature shift to avoid early lethality, is sufficient to decrease *bru1* mRNA levels in IFMs (Fig. S5 C), supporting that Rbfox1 can indeed negatively regulate Bru1 levels. This indicates that in IFMs, the expression level of Rbfox1 is tightly regulated and determines if Rbfox1 negatively or positively influences Bru1 expression. This regulation is likely fiber-type specific, as *bru1* mRNA levels in *Rbfox1-*IR^27286^ TDT are decreased (Fig. S5 D) and protein levels of both Bru1-PA and Bru1-PB tend to increase in *Rbfox1-*IR^27286^ TDT and Abd (Fig. S5 E, F). Similar fiber-type and level-dependent regulation were also observed for *exd* and *salm* mRNAs, as discussed below.

We next evaluated if Bru1 might regulate Rbfox1. *Rbfox1* mRNA levels are significantly downregulated in mRNA-Seq data from 72h APF pupae and 1d adults (Spletter et al., 2015) when *bru1* is knocked down in IFMs using RNAi (Fig. S5 G), suggesting that Bru1 positively regulates Rbfox1 expression. However, there is no significant effect on *Rbfox1* mRNA levels in IFMs or TDT from *bru1^M2^* or *bru1^M3^* mutants (Fig. S5 G, I, J), suggesting this regulation depends on how much Bru1 protein is actually present in the muscle. We see a similar effect when Bru1 is overexpressed: early and strong overexpression in IFMs with the Mef2 driver significantly decreases *Rbfox1* mRNA levels (Fig. S5 H), but overexpression from 34h APF with UH3-Gal4 (IFM) does not (Fig. S5 I). Overexpression of Bru1 in TDT with Act79B-Gal4 also tends to reduce *Rbfox1* levels, although this was not statistically significant (Fig. S5 J), suggesting that Bru1 can also negatively regulate *Rbfox1* mRNA levels. We conclude that Bru1 can regulate Rbfox1 levels in *Drosophila* muscle, and likely in a level-dependent manner.

### Rbfox1 and Bru1 genetically interact selectively during IFM development

Having established that Rbfox1 and Bru1 regulate each other’s expression, we next explored if they might cooperatively regulate muscle development. *bru1*-IR is reported to result in short sarcomeres and hypercontraction (Oas et al., 2014; Spletter et al., 2015), a phenotype very similar to what we characterized in *Rbfox1* knockdown (Fig. 3). We verified that the Bru1 phenotype is IFM-specific in *bru1^M2^* mutants (Fig. 6) and *bru1*-IR flies (Fig. S6). We observed loss of myofibers (Fig. 6 B, Fig. S6 C) as well as short, thick sarcomeres in the IFMs (Fig. 6 F, Q, R, Fig. S6 G), but no phenotype in either TDT or Abd-M (Fig. 6J, N, S, T, Fig. S6 K, O). By contrast, *Rbfox1* knockdown affects tubular as well as fibrillar muscles (Fig. 6 C, G, K, O; Fig. S6 B, F, J, N). To test if overexpression of Bru1 can also induce hypercontraction like we observe with overexpression of Rbfox1 (Fig. S2 E), we drove UAS-Bru1 using Mhc-Gal4 (which expresses from 40h APF onwards). Indeed, overexpression of Bru1 leads to an IFM hypercontraction phenotype including myofiber loss (Fig. S6 R) and torn myofibrils with short sarcomeres (Fig. S6 R’). This phenotype could be partially rescued by the *Mhc^P401S^* allele of myosin heavy chain (Fig. S6 S, S’), confirming that myofiber detachment is indeed due to hypercontraction. Thus, loss as well as gain of both Bru1 and Rbfox1 in IFM result in similar phenotypes, including hypercontraction.

**Figure 6:**
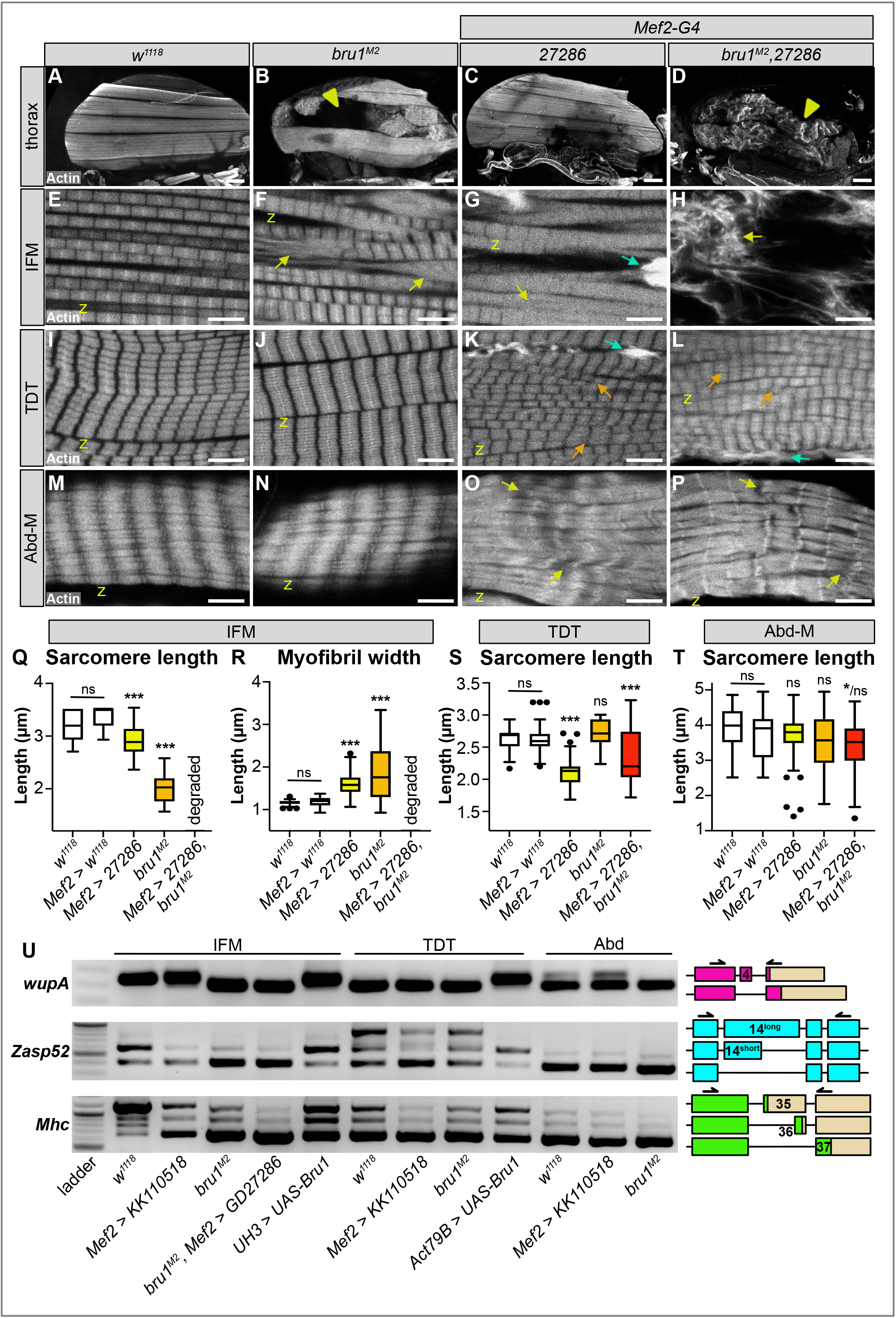
Rbfox1 and Bru1 genetically interact in IFM myogenesis and regulate the alternative splicing of sarcomere genes. **A-D)** Confocal projections of hemithoraces showing IFMs (A-D) from *w^1118^*, *bru1^M2^*, *Rbfox1-IR^27286^* and *bru1^M2^*, *Rbfox1-IR^27286^* flies. Arrowheads indicate aberrant, torn myofibers. Scale bars = 100 μm. **E-H)** Single-plane confocal images from IFM, showing torn myofibrils (yellow arrows) with short sarcomeres and actin inclusions (cyan arrows) in *bru1^M2^* (F) and *Rbfox1-IR^27286^* (G). *bru1^M2^*, *Rbfox1-IR^27286^* demonstrates genetic interaction and loss of myofibril structure (H). **I-P)** Single-plane confocal images from TDT (I-L) and Abd-M (M-P) from *w^1118^*, *bru1^M2^*, *Rbfox1-IR^27286^* and *bru1^M2^*, *Rbfox1-IR^27286^* flies. Myofibrils in *Rbfox1* knockdown muscles are disorganized (orange arrows), have actin inclusions (cyan arrows) and are often torn (yellow arrows). Scale bars = 5 μm. **Q-R)** Quantification of sarcomere length (Q) and myofibril width (R) in IFM. **S-T)** Quantification of sarcomere length in TDT (S) and Abd-M (T). Significance determined in comparison to *w^1118^* by ANOVA and post-hoc Tukey (not significant, ns; *= p < 0.05; *** = p < 0.001). **U)** RT-PCR for select alternative splice events in *wupA* (magenta), *Zasp52* (blue) and *Mhc* (green). Genotypes as labeled. Primer locations and alternative isoforms are diagrammed on the right. Exon numbers are based on annotation FB2021_01. UTR regions, tan.

This led us to test what happens to muscles lacking both Rbfox1 and Bru1. Knockdown with *Rbfox1-*IR^27286^ in the *bru1^M2^* background reveals a strong genetic interaction. IFM myofibers were still present but severely disorganized and displayed an unusual banded actin pattern (Fig. 6 D). Myofibril and sarcomere structure were completely compromised and F-actin formed into disarrayed clumps, as well as spine and star-like structures (Fig. 6 H). We obtained an identical IFM phenotype in double knockdown (*bru-IR*, *Rbfox1*-RNAi) flies with Mef2-Gal4 expression restricted to adult IFM development using *Tubulin-Gal80^ts^* and a temperature shift at the late third instar larval stage (Fig. S6 D, H). This genetic interaction is restricted to IFMs, as the phenotype in TDT and Abd-M was not enhanced and appeared consistent with the phenotype observed in *Rbfox1-*IR^27286^ (compare Fig. 6 K, O to L, P) or *Rbfox1*-RNAi (compare Fig. S6 J, N to L, P) alone. TDT myofibrils were disorganized and frayed with short sarcomeres (Fig. 6 L, S; Fig. S6 L), while Abd-M myofibrils were discontinuous and sarcomere structure was irregular (Fig. 6 P, T; Fig. S6 P). This result indicates that Rbfox1 and Bru1 genetically interact in fibrillar IFM, but not in tubular TDT and Abd-M where primarily Rbfox1 seems to function.

### Rbfox1 and Bruno1 co-regulate alternative splice events in IFMs

Many developmentally-regulated, alternatively spliced exons in vertebrate muscle have binding sites for both FOX and CELF family RNA-binding proteins, and in heart notably appear to be antagonistically co-regulated by CELF2 and RBFOX2 (Bland et al., 2010; Gazzara et al., 2017). Thus, we next checked if Rbfox1 and Bru1 co-regulate alternative splicing in *Drosophila* muscle. We performed RT-PCR for select alternative splice events in structural proteins known to have fibrillar and tubular specific isoforms, including *TnI (wupA)*, *Zasp52*, *Mhc*, *Sls* and *Strn-Mlck*. TnI has IFM- and TDT-specific protein isoforms marked by the presence or absence of exon-4 (based on the most recent Flybase annotation, formerly exon 3) (Fig. S6 Q) (Barbas et al., 1993; Beall and Fyrberg, 1991), and this splice event is regulated by Bru1 (Oas et al., 2014; Spletter et al., 2015). Loss of *bru1* but not *Rbfox1* in IFMs caused a complete switch to the tubular event promoting *wupA-Ex4* skipping (Fig. 6 U). Overexpression of Bru1 with Act79B-Gal4 in TDT was sufficient to switch to the IFM event and restore splicing into exon-4 (Fig. 6 U). RT-PCR selective for *wupA-Ex4* revealed an overall decreased expression in *Rbfox1* knockdown IFMs and TDT, and complete loss in *bru1^M2^* mutant muscle (Fig. S6 T). These results suggest that *wupA-Ex4* splicing is largely dependent on Bru1, and changes to *wupA* splicing in *Rbfox1* knockdown are likely indirect.

Another structural protein with fibrillar and tubular specific isoforms known to be regulated by Bru1 is Zasp52 (Spletter et al., 2015). *Zasp52* exon-14 is preferentially included in TDT, shortened in IFMs and skipped in Abd-M (Fig. 6 U). In IFMs, knockdown of *Rbfox1* and loss of Bru1 results in a shift towards exon-14 skipping. In TDT, knockdown of *Rbfox1* also results in a shift towards exon skipping, while loss of Bru1 has little effect. Overexpression of Bru1 in TDT is sufficient to shift splicing to the “short exon 14” isoform of *Zasp52*. Neither *Rbfox1* knockdown nor loss of Bru1 alters *Zasp52* splicing in Abd. This result indicates that Bru1 promotes use of the alternative 3’ splice site leading to a “short exon 14” *Zasp52* isoform in both TDT and IFMs. In TDT but not in IFMs or Abd, Rbfox1 promotes inclusion of full-length exon 14, independent of Bru1.

Myosin Heavy Chain (Mhc) has three different alternative C-terminal exons that are differentially spliced in a temporal and muscle-type specific manner (Clyne et al., 2003; Kao et al., 2019; Orfanos and Sparrow, 2013). The IFMs in adult flies preferentially use the first termination site encoded by exon 35 (Fig. 6 U). In *Rbfox1* knockdown and *bru1^M2^* mutant IFMs, there is a shift towards use of the third termination site in exon 37, and this shift is more accentuated in *Rbfox1-IR^27286^*, *bru1^M2^* IFMs (Fig. 6 U), suggesting that both Rbfox1 and Bru1 control this event. Use of all three terminal exons is detected in TDT, although exon 37 is preferential. *Rbfox1-IR^KK110501^* TDT shows a shift towards almost exclusive use of exon 37 (Fig. 6 U). There is little or no effect on *Mhc* splicing in TDT from *bru1^M2^* mutants or with Bru1 overexpression. Exon 37 is preferentially used in Abd, and loss of Rbfox1 and Bru1 has little effect on *Mhc* splicing (Fig. 6 U). This suggests that both Rbfox1 and Bru1 control *Mhc* C-terminal splicing in IFM, but predominantly Rbfox1 directs *Mhc* splicing in TDT.

We also tested two additional fiber-type specific splice events in *Strn-Mlck* and *Sls*. The Strn-Mlck Isoform R protein, produced from a transcript containing an early termination in exon 25, is specifically expressed in IFMs (Spletter et al., 2015), although by RT-PCR we could amplify the *Strn-Mlck-RR* mRNA in all muscle types (Fig. S6 U). This event is dependent on Bru1 and independent of Rbfox1 in IFM, TDT and Abd (Fig. S6 U). Sls exon 10 was previously shown to be included in tubular muscle but skipped in IFMs in a Bru1-dependent manner (Oas et al., 2014). We confirmed that the *sls* isoform skipping exon 10 is absent in *bru^M2^* mutant IFMs, TDT and Abd-M, and Bru1 overexpression is sufficient to promote skipping in IFMs and TDT (Fig. S6 U). In *Rbfox1-IR^KK110501^* flies, we observed a slight decrease in exon 10 skipping in IFMs, no change in TDT and a slight increase in exon 10 skipping in Abd. Taken together, our data suggest a complex regulatory dynamic where Rbfox1 and Bru1 co-regulate some alternative splice events and independently regulate other events in a muscle-type specific manner.

### Rbfox1 regulates myofiber fate determining transcriptional activators

Fiber-type identity and muscle type-specific gene expression is both specified and maintained through transcriptional regulation (Spletter and Schnorrer, 2014). Our bioinformatic analysis identified Rbfox1 binding motifs in more than 40% of transcription factors genes (Figure S3 C), notably including *Mef2*, *extradenticle* (*exd*), *homothorax* (*hth*), *E2F transcription factor 1* (*E2f1*), *DP transcription factor* (*Dp*), *apterous* (*ap*)*, twist* (*twi*), *cut* (*ct*), *vestigial* (*vg*) and *scalloped* (*sd*) (Table S1), which have all been shown to regulate adult muscle identity or myofiber gene expression (Dobi et al., 2015; Zappia and Frolov, 2016). Interestingly, even though it lacks Rbfox1 binding motifs, we observed regulation of Act88F expression in *Rbfox1*-RNAi IFMs. Thus, we next tested if Rbfox1 regulates transcriptional activators which could in turn regulate structural gene expression.

We first evaluated expression of *extradenticle* (*exd*), a gene encoding a homeodomain protein which is suggested to be genetically upstream of Salm and Bru1 and in particular was shown to direct expression of *Act88F* (Bryantsev et al., 2012b). *exd* contains three Rbfox1 binding sites, one in an intron and two in the 3’-UTR (Fig. S3 F). *exd* transcript levels were significantly down-regulated in IFMs from *Rbfox1-IR^KK110518^* knockdown flies (Fig. 7 A). This regulation is likely Rbfox1 level-dependent, as weaker knockdown with both *Rbfox1*-RNAi and *Rbfox1-IR^27286^* tended towards increased *exd* levels in IFM (Fig. 7A). We were not able to detect Rbfox1 binding to *exd* mRNA in RIP from adult thoraces of *Rbfox1^CC00511^* flies (data not shown), but we cannot rule out binding at earlier stages of muscle development. This indicates that Rbfox1 can regulate *exd* levels, but the nature of this regulation requires further investigation.

**Figure 7:**
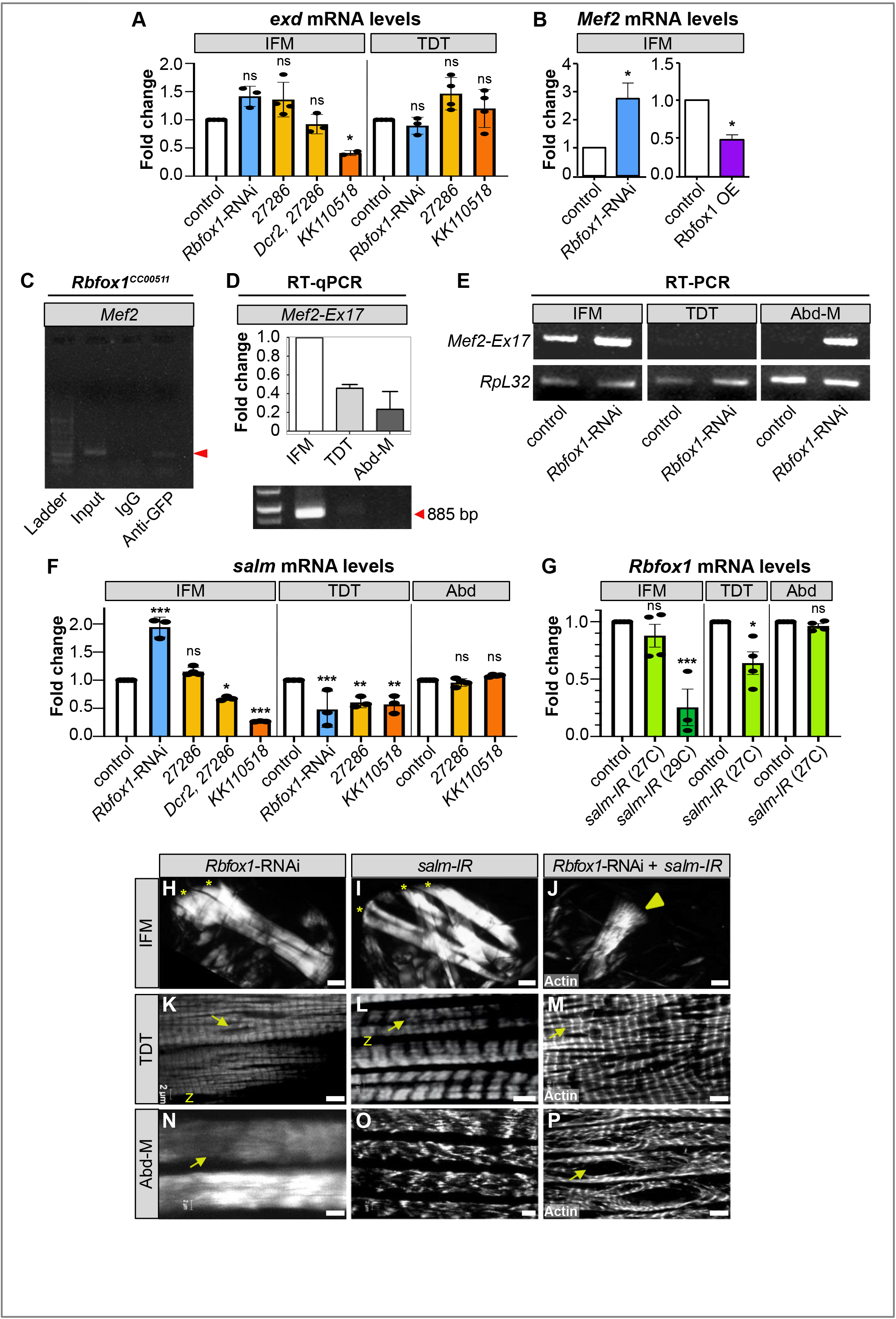
Rbfox1 regulates expression of myogenic transcription factors *exd* and *Mef2* and genetically interacts with *salm* in IFM development. **A)** RT-qPCR (*Rbfox1*-RNAi) and semi-quantitative RT-PCR (*Rbfox1-IR^27286^*, *Rbfox1-IR^KK110518^*) quantification of the fold change in *exd* transcript levels in IFM and TDT across the *Rbfox1* knockdown series. Data was normalized by *RpL32* levels. **B)** RT-qPCR quantification of the fold change in *Mef2* mRNA expression in IFM with *Rbfox1*-RNAi or Rbfox1 OE. Significance is from paired t-test (* = p < 0.05). **C)** RIP using the *Rbfox1^CC00511^* line followed by RT-PCR indicates Rbfox1 binds to *Mef2* mRNA (red arrowhead). **D)** Semi-quantitative RT-PCR demonstrating that *Mef2* isoforms containing exon 17 and thus a short 5’-UTR (see also Fig. S3 G, Fig. S7 A, B) are preferentially expressed in wildtype IFM. **E)** RT-PCR detects increased use of *Mef2-Ex17* in *Rbfox1*-RNAi IFM and Abd-M. **F)** RT-qPCR (*Rbfox1*-RNAi) and semi-quantitative RT-PCR (*Rbfox1-IR^27286^*, *Rbfox1-IR^KK110518^*) quantification of the fold change in *salm* transcript levels in IFM, TDT and Abd across the *Rbfox1* knockdown series. Data was normalized by *RpL32* levels. **G)** Fold change in *Rbfox1* transcript levels in IFM, TDT and Abd normalized to *RpL32* after *salm-IR* at 27 °C or 29 °C, as determined by RT-qPCR (29 °C) and semi-quantitative RT-PCR (27 °C). Significance in A, F, G determined by ANOVA and post-hoc Tukey (not significant, ns; *= p < 0.05, **= p < 0.01, ***= p < 0.001), error bars indicated standard deviation. **H-J)** Polarized microscopy images of hemithoraces showing a reduction in myofiber number (stars) with *Rbfox1*-RNAi (H) and *salm-IR* (I), and a complete loss of IFMs with double *Rbfox1*-RNAi, *salm-IR* knockdown (J). TDT, yellow arrowhead; quantification, Fig. S7 K). Scale bars = 100 μm. **K-P)** Single plane confocal images of TDT (K-M) and Abd-M (N-P) showing abnormal myofibril structure and tearing (arrows) in *Rbfox1*-RNAi, *salm-IR*, and *Rbfox1*-RNAi, *salm-IR* knockdown tubular muscle. Quantification, Fig. S7 K; Scale bars = 5 μm.

We next evaluated expression of *Mef2*, a well-characterized MADS-box transcription factor that regulates structural protein expression (Molkentin et al., 1995; Tanaka et al., 2008). *Mef2* contains five intronic, one 5’-UTR and one 3’-UTR Rbfox1 binding motifs (Fig. S3 G). *Mef2* mRNA levels were significantly up-regulated in IFMs with *Rbfox1*-RNAi and significantly down-regulated with Rbfox1 over-expression (Fig. 7 B). We were able to detect Rbfox1 binding to *Mef2* mRNA in RIP from adult thoraces of *Rbfox1^CC00511^* flies (Fig. 7 C), suggesting this regulation may be direct. As Rbfox1 binding sites in *Mef2* are concentrated in the upstream introns, we wondered if they might influence alternative 5’-UTR use. In our mRNA-Seq data, we observed both temporal and fiber-type specific use of *Mef2* 5’-UTR exons. The short 5’-UTR encoded by *Mef2-Ex17* is preferential to developing IFMs (Fig. S7 A, B), which we could confirm using qPCR (Fig. 7 D). The longer 5’-UTR encoded by *Mef2-Ex20* is used in all muscles as they mature, while a second long 5’-UTR encoded by *Mef2-Ex21* is predominant in developing tubular muscle and myoblasts (Fig. S7 A, B). Interestingly, we could detect increased use of *Mef2-Ex17* in IFMs and Abd-M from adult *Rbfox1-*RNAi flies (Fig. 7 E) and a trend towards increased use of *Mef2-Ex20* and *Mef2-Ex21* in IFMs from *bru1-IR* and *bru1^M3^* flies (Fig. S7 A), suggesting that Rbfox1 and Bru1 may regulate use of these variable *Mef2* 5’-UTR regions.

Levels of Mef2 are known to affect muscle morphogenesis but not production of different isoforms (Gunthorpe et al., 1999), thus we next examined whether increased Mef2 levels can induce muscle hypercontraction. Although Mef2-Gal4 driven overexpression of UAS-*Mef2* caused lethality after 48 hours, flies with Mhc-Gal4 driven overexpression survive to adulthood. These flies were flightless, displayed sarcomeric defects (Fig. S7 C, C’) and had increased levels of TnI and Act88F in IFMs (Fig. S7 D, E). Notably, they do not display a hypercontraction defect. We conclude that increased levels of Mef2 can lead to an overall increase in many structural proteins, but hypercontraction observed upon changes in Rbfox1 and Bru1 levels likely results from alternative splicing defects and a possible isoform-imbalance amongst the structural proteins.

As a third and final example, we investigated if Rbfox1 regulates Spalt major (Salm), a C2H2-type zinc finger transcription factor that serves as master regulator of the fibrillar muscle fate (Schönbauer et al., 2011a). Although *Salm* does not contain canonical Rbfox1 binding motifs, its expression is controlled by the homeodomain proteins Extradenticle (Exd) and Homothorax (Hth) (Bryantsev et al., 2012b) as well as Vestigial (Vg) and its co-factor Scalloped (Sd) (Schönbauer et al., 2011a). Salm is speculated to influence muscle diversification by modification of Mef2 level (Spletter and Schnorrer, 2014), and is known to regulate expression of *bru1*, *wupA (TnI)* and *Act88F* (Schönbauer et al., 2011a; Spletter et al., 2015; Spletter et al., 2018). Thus, we wanted to determine if it might interact with the Rbfox1 regulatory hierarchy.

We first examined *Salm* mRNA levels in Rbfox1 knockdown muscle. *Salm* levels were significantly increased in IFM from *Rbfox1-*RNAi animals, but significantly decreased in IFMs from flies with Dcr2 enhanced *Rbfox1-IR^27286^* or *Rbfox1-IR^KK110518^* (Fig. 7 F). *Salm* levels in TDT were significantly decreased in all knockdown conditions, and were not affected in Abd (Fig. 7 F). This suggests that Rbfox1 can regulated *Salm*, and in the IFMs this regulation is dependent on the level of Rbfox1 expression. We also could show that *Rbfox1* mRNA levels were significantly decreased in both IFMs and TDT, but not in Abd, of *Salm-IR* flies (Fig. 7 G). As Salm is the master regulator of the fibrillar muscle fate, these results suggest there is cross-regulation between identity transcription factors and fiber-type specific splicing networks.

To investigate the physiological relevance of this interaction, we knocked down both *Salm* and *Rbfox1* in all muscle fiber types using Mef2-Gal4. We first confirmed that *Salm-IR* is efficient (Fig. S7 I) and verified previous findings (Schönbauer et al., 2011a) that *Salm-IR* results in a tubular-muscle fate conversion of the IFMs and a loss of *bru1* expression (Fig. 7 I, Fig. S7 H, J). We also observed mild defects in myofibrillar patterning in the TDT with both *Salm-IR* (Fig. 7 L) and in *FRT-salm-FRT* mutants (Fig. S7 F-G). Double knockdown with *Rbfox1*-RNAi and *Salm-IR* resulted in greater than 60% lethality and severe locomotion defects (data not shown). IFMs were completely missing in hemi-thoraces from double knockdown flies (Fig. 7 J, quantification in Fig. S7 K), and although TDT was present, both myofibril structure and organization were aberrant (Fig. 7 M). Abd-M were also disorganized and frequently torn (Fig. 7 P, Fig. S7 K). This data indicates there is indeed a genetic interaction between Salm and Rbfox1 in IFM- and TDT-development that is necessary for proper fiber-type gene expression and alternative splicing. Altogether, our results suggest that Rbfox1 is involved in the regulation of fiber specific isoforms of structural proteins, particularly TnI, not only through directly regulating the splicing process, but also through hierarchical regulation of the fiber diversity pathway.

## Discussion

Here we report the first detailed characterization of Rbfox1 function in *Drosophila* muscle. We show that Rbfox1 functions in a fiber-type and level-dependent manner to modulate both fibrillar and tubular muscle development. Collectively, our data demonstrate that Rbfox1 operates in a complex regulatory network to fine-tune the transcript levels and alternative splicing pattern of fiber-type specific structural proteins such as Act88F, TnI, Strn-Mlck, Zasp52 and Mhc (Fig. 8 A). It does this directly, by binding to 5’-UTR and 3’-UTR regions to regulate transcript levels and binding to intronic regions to promote or inhibit alternative splice events. In addition, Rbfox1 regulates transcriptional activators and other splicing factors such as Bru1 which themselves regulate transcript levels and alternative splicing events (Fig. 8 A). We found the valence of several regulatory interactions for both Rbfox1 and Bru1 to be expression level dependent in IFM (Fig. 8 B), suggesting this regulatory network is carefully balanced to respond to even small changes in gene expression. Moreover, as in vertebrates, Rbfox1 and Bru1 exhibit cross-regulatory interactions (Fig. 8 B) and genetically interact in IFM development. Interestingly, this cross-regulation extends to Salm, which suggests that RBPs such as Rbfox1 might actively regulate transcriptional networks to guide and refine acquisition of fiber-type specific properties during muscle differentiation.

**Figure 8:**
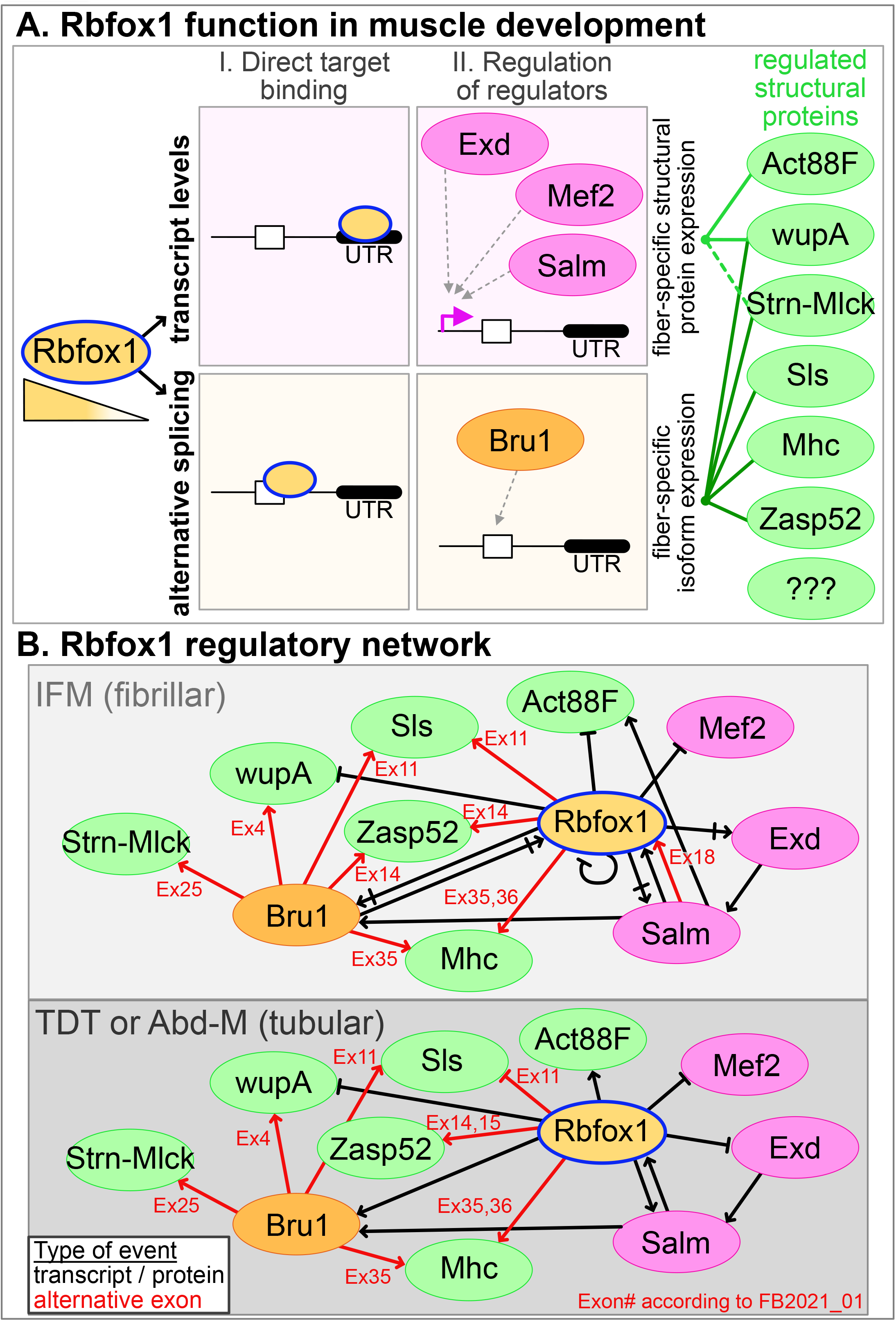
Model of the Rbfox1 fiber-type specific regulatory network and function in *Drosophila* muscle development. **A)** Rbfox1 regulates transcript levels and alternative splicing of target genes. It can do this directly by binding UTR or intronic sequences, respectively. Rbfox1 also regulates levels of transcriptional activators such as Exd, Mef2 and Salm which in turn affect transcription levels as well as the RNA-binding protein Bru1 which regulates alternative splicing. Ultimately, this defines fiber-type specific expression levels and splice isoform usage of sarcomeric genes. RNA-binding proteins, orange; Rbfox1, blue outline; transcription factors, magenta; structural proteins, green. **B)** Rbfox1 regulatory network events confirmed in this manuscript in fibrillar IFM (light grey, top) or tubular TDT or Abd-M (dark grey, bottom). Symbol definitions: arrow, positive regulation; flat-ended arrow, negative regulation; double-ended arrow, level-dependent bivalent regulation; circular arrow, autoregulation; paired arrows, cross-regulation; red, alternative exon; black, transcript or protein level. Oval fill colors as in A. Exon numbers (Ex) according to annotation release FB2021_01.

### Rbfox1 function in muscle development is evolutionarily conserved

Although the conserved nature of the Rbfox1 binding site (5’-UGCAUG-3’) in mammalian genomes is well known (Denisov and Gelfand, 2003; Jin et al., 2003), its functional significance in muscle was appreciated only after genome wide studies showing that the regulatory element is enriched in introns flanking skeletal and cardiac muscle specific exons in humans and mice (Castle et al., 2008; Kalsotra et al., 2008). Transcripts of hundreds of structural genes are mis-spliced in Rbfox1 and Rbfox2 knockout mice, which developmentally have defects in muscle structure and function, and as adults fail to maintain skeletal muscle mass (Pedrotti et al., 2015; Singh et al., 2018). Knockdown of Rbfox1 and Rbfox2 in zebrafish leads to defects in alternative splicing, myofiber morphology, and function of both heart and skeletal muscle (Gallagher et al., 2011). Even mutants in the *C. elegans* homologue *fox-1* lead to aberrant myoblast migration and impaired egg-laying (Kuroyanagi et al., 2006; Mackereth, 2014). We previously reported that muscle-specific knockdown of *Rbfox1* in *Drosophila* results in short IFM sarcomeres (Nikonova et al., 2019). Here we extend those findings and show that as in vertebrates, Rbfox1 binding sites in the *Drosophila* genome are enriched in the introns and UTR-regions of muscle genes. Rbfox1 knockdown affects all adult muscle fiber-types and is characterized by defects in muscle-specific alternative splicing, myofibril and sarcomere structure and impaired muscle function. As tubular muscles in fly reflect the vertebrate skeletal muscle physiology and IFMs share characteristics with cardiac muscle, our observations are consistent with observations in vertebrates and strongly suggest that the function of Rbfox1 in muscle development is evolutionarily conserved.

### Rbfox1 regulates fiber-type specific isoform switches during development

Studies from both vertebrates and *C. elegans* suggest that Rbfox1 modulates developmental isoform switches. In mouse, Rbfox1 and Rbfox2 regulate splicing of *Mef2D exon α2* during myotube differentiation allowing Mef2D to escape inhibitory PKD signaling and activate the late-muscle gene expression program (Runfola et al., 2015). In *C. elegans*, FOX-1/ASD-1 and SUP-12 regulate a developmental switch in expression of the fibroblast growth factor receptor *egl-15* that is necessary for myoblast migration and vulval muscle formation (Kuroyanagi et al., 2007; Mackereth, 2014). Rbfox1 is upregulated as cardiac cells differentiate and knockdown results in cardiac hypertrophy and splicing defects, consistent with the reduction in Rbfox1 expression in human patients with dilated cardiomyopathy and in hypertrophic heart from mouse and zebrafish (Gao et al., 2016). In myotonic dystrophy (DM1), where muscle exhibits a reversion from mature to embryonic splicing patterns (Blech-Hermoni et al., 2016), dystrophic muscle in mouse models and human patient cells produces a dominant-negative Rbfox1 isoform through mis-regulated alternative splicing that enhances DM1 phenotypes (Klinck et al., 2014). Previous studies in *Drosophila* have also identified transcriptional and isoform switches during normal IFM development (Burkart et al., 2007; González-Morales et al., 2019; Nongthomba et al., 2007; Orfanos and Sparrow, 2013; Spletter et al., 2018), and indeed we find that not only are Rbfox1 levels temporally regulated in IFM, but also splicing of genes with characterized isoform switches including *sls*, *Mhc, zasp52* and *wupA* is altered after Rbfox1 knockdown. Notably, most of these events are also muscle fiber-type specific and result in the production of fibrillar and tubular specific isoforms (Spletter et al., 2015; Venables et al., 2012). This implies the developmental function of Rbfox1-mediated splicing is to establish fiber-type specific properties during muscle differentiation. Our data suggests it would be informative to investigate differences in Rbfox1 function between fiber-types in vertebrate models, as the role of Rbfox1 in generating fiber diversity is likely to be conserved in higher vertebrates and disease-relevant.

### Rbfox1-mediated splicing is subject to cross-regulatory interaction with Bru1

The interactions between RBPs are important in defining alternative splicing patterns in muscle. For example, hnRNP-G and Tra2*β*, which are predominant in cardiac and skeletal myoblasts, respectively, have opposing effects on splicing of a *Dystrophin* exon that is abnormally incorporated in heart muscles of human patients with X-linked dilated cardiomyopathy (Nasim et al., 2003). Rbfox binding motifs are found to be co-enriched with MBNL and CELF motifs around the same groups of exons in human, mouse and chicken (Bland et al., 2010; Kalsotra et al., 2008; Merkin et al., 2012). Rbfox1 and MBNL co-regulate a significant number of alternative events altered in DM1 skeletal muscle (Klinck et al., 2014), while CELF2 and Rbfox2 co-regulate and co-bind introns flanking exons regulated in cardiac development or with altered expression in hearts of a Type I diabetes mouse model (Gazzara et al., 2017). CELF2 moreover represses Rbfox2 expression in heart, and overexpression of CELF1/2 or depletion of Rbfox2 leads to the same changes in splicing direction and magnitude (Gazzara et al., 2017). We see evidence for similar regulatory interactions between Rbfox1 and the CELF1/2 homolog Bru1 in our data from *Drosophila*. Loss of either Rbfox1 or Bru1 can lead to muscle hypercontraction, a condition similar to myopathies seen in reperfused rat hearts (Duncan, 1987; Monticello et al., 1996). Rbfox1 and Bru1 cross-regulate each other’s expression and co-regulate alternative splicing of events in *Mhc*, *zasp52*, *sls*, and *wupA*. Our data provide novel insight into this regulatory interaction, as we show the valence of both cross-regulation and splicing events is expression-level and fiber-type dependent. Moreover, Rbfox1 and Bru1 genetically interact in IFM development as knockdown of both RBPs leads to complete loss of myofibril structure. Our data show that *Drosophila* is an informative model for future studies to unravel conserved, fiber-type specific mechanisms of RBP cross-regulation, cooperation, antagonism and feedback on a genome-wide scale.

### Rbfox1 modulates fiber-type specific transcriptional networks

Although it is an RBP, Rbfox is reported to modulate transcriptional networks. Rbfox2 can interact with the Polycomb repressive complex 2 (PRC2) through a unique C-terminal domain and regulate transcription in mouse (Wei et al., 2016). In *Drosophila*, Rbfox1 can interact with Cubitus interruptus (Ci) and Suppressor of Hairless (Su(H)), transcription factors in the Hedgehog (Hh) and Notch (N) signaling pathways, respectively, to regulate vein-intervein and sensory organ specification in the wing disc (Shukla et al., 2017; Usha and Shashidhara, 2010). Our data indicate that in fly muscle, in contrast to these examples of protein-level interaction, Rbfox1 regulates mRNA transcript levels of transcription factors including *exd*, *salm*, and *Mef2*. Although the mechanism of *salm* regulation is unclear, *exd* and *Mef2* both are potentially regulated by direct 3’-UTR binding and/or through the splicing of alternative 5’-UTR sequences. It remains to be tested if the short 5’-UTR of *Mef2* negatively regulated by Rbfox1 is more or less stable than the long 5’-UTR preferentially used in tubular muscle. Interestingly, Rbfox1 regulates splicing of a MEF2A exon in mouse and zebrafish heart that is mis-spliced in cells from human patients with dilated cardiomyopathy (Gao et al., 2016), and Rbfox1 and Rbfox2 cooperatively regulate splicing of Mef2D during C2C12 differentiation (Runfola et al., 2015). Our data thus support findings that Rbfox1 modulates transcription, but introduce a novel method of regulation, via regulating transcription factor transcript stability.

The conserved regulation of Mef2 by Rbfox proteins is particularly intriguing, as Mef2 is a key regulator of expression of most structural proteins during assembly of the sarcomere (Gunthorpe et al., 1999; Molkentin et al., 1995; Tanaka et al., 2008; Taylor and Hughes, 2017). In *Drosophila*, differential expression levels of Mef2 define corresponding fiber-type specific expression levels of structural proteins (Gunthorpe et al., 1999; Hughes et al., 1993). Given the thin to thick filament ratio is 6:1 in fibrillar muscles, and 8-12:1 in the tubular muscles (Bernstein et al., 1993), a fiber specific isoform of Mef2 might explain fiber specific changes in the expression of sarcomeric proteins. Additionally, knockdown of *Rbfox1* is able to partially rescue the hypercontraction phenotype in *wupA^hdp-3^* heterozygotes, indicating a role for Rbfox1 in maintaining the stoichiometry of structural proteins by regulating splicing/expression of TnI. Increased expression of Mef2, Bru1 and Salm combined with inclusion of IFM-specific *wupA-Ex4*, all favoured by low levels of Rbfox1, could generate a stoichiometric imbalance resulting in hypercontraction in the *Rbfox1* knockdown condition.

The fibrillar muscle fate is specified transcriptionally, where expression of Vestigial (Vg), Extradenticle (Exd) and Homothorax (Hth) in IFM progenitors induces Salm expression (Bryantsev et al., 2012b; Schönbauer et al., 2011b). Salm further instructs the fibrillar fate by directly or indirectly inducing Bru1 and more than 100 fibrillar-specific genes (Oas et al., 2014; Spletter et al., 2015). The mammalian ortholog of Salm, Sall1, is also involved in fate determination of cardiomyoblasts in mice (Morita et al., 2016). Studies so far report the positive regulation of these factors, but here we report the first evidence for negative regulation for fine tuning acquisition of muscle-type specific properties. Depending on its expression level, Rbfox1 can either promote or inhibit expression of *exd*, *salm* and *bru1*. Notably, Rbfox1 promotes expression of the *bru1-RB* isoform which is preferentially used in TDT, indicating Bru1 might have isoform-specific functions in different fiber-types. This is also possible for Rbfox1 itself, as Rbfox1 nuclear and cytoplasmic isoforms are reported to have distinct functions (Hamada et al., 2016) and we observe fiber-type differential use in Rbfox1 exons. In addition, we show that Salm positively regulates Rbfox1 levels in both IFM and TDT. This multi-level, cross-regulatory loop suggests that the fiber diversification network continuously integrates both RBP and transcriptional feedback to refine expression levels of key regulatory components, here Bru1, Rbfox1, Salm, Exd and Mef2, to ultimately fine-tune the expression level and ratio of structural protein isoforms. Such a mechanism may be broadly applicable to allow muscle fibers to flexibly adjust regulator levels during development, or to promote plasticity in response to exercise, aging, injury or disease.

## Materials and Methods

A table of key resources is available as Supplemental Table 3.

### Fly stocks and crosses

Fly stocks were maintained using standard culture conditions. Wildtype controls include either *w^1118^ or Canton-S*. Rbfox1-GFP (*Rbfox1^CC00511^*) was generated as part of a protein enhancer trap library (Kelso et al., 2004), and does not alter protein function or localization. Fly stocks of UAS-*Rbfox1*-RNAi and UAS-*Rbfox1* were kind gifts from L. Shashidhara, IISER, Pune, India (Usha and Shashidhara, 2010). The deGrad-FP fly line *pUASP1-deGradFP/CyO; MKRS/TM6,Tb* (Caussinus et al., 2012) was a kind gift of Sonal Jaishwal, CCMB, India. deGradFP knockdown was carried-out during adult IFM development by temperature shifts of late third instar larvae. *Mhc^P401S^* (Nongthomba et al., 2003) is a myosin mutant that minimizes acto-myosin force in IFM, while *wupA^fliH^* (Firdaus et al., 2015) and *wupA^hdp3^* (Barbas et al., 1993) are known hypercontraction mutants in *wupA* (TnI). RNAi lines were obtained from the Vienna Drosophila Resource Center (VDRC) including UAS-*Arrest*-RNAi (*Bru1-IR)* (41547, 48237, 41568) (Dietzl et al., 2007; Oas et al., 2014; Spletter et al., 2015), UAS-Salm-RNAi (*salm-IR*) (3029, 101052) (Schönbauer et al., 2011a), UAS-*Rbfox1*-IR^KK110518^ (110518) or from the Bloomington Drosophila Stock Center (BDSC) UAS-*Rbfox1*-IR^27286^ (TRiP27286). UAS-Mef2 lines were provided by Alberto Ferrus (Gunthorpe et al., 1999). UAS-Bru1-PA (also called UAS-*Arrest*) was kindly provided by Richard Cripps (Oas et al., 2014) and expresses the full-length *bru1-RA* mRNA from DGRC clone LD29068. A second UAS-Bru1-PA line was generated by cloning the full-length *bru1-RA* cDNA (obtained by RT-PCR from *w^1118^*) into the *pUAS-TattB* transformation vector (Bischof et al., 2007), and integrating into the attP-86Fb landing site. The *bru1^M2^* and *bru1^M3^* deletion alleles were generated using a CRISPR approach as described in (Zhang et al., 2014), where the C-terminal portion of the *bru1* coding region (including RRM2, the divergent domain and RRM3 for *bru1^M2^* and RRM3 and the 3’-UTR for *bru1^M3^*) was replaced by a selectable 3xP3-DsRed cassette. sgRNA sequences and homology arm primers are listed in Supplemental Table 2. Gal4 drivers used were: Mef2-Gal4 (Ranganayakulu et al., 1996) which drives in all muscle (maintained at 27 °C or 29 °C); UAS-Dcr2, Mef2-Gal4 which enhances RNAi efficiency (maintained at 22°C); Act5c-Gal4 which drives in all cells (maintained at 27°C and 25°C); Mhc-Gal4 (Davis et al., 1996); UH3-Gal4 (Singh et al., 2014) is a driver with IFM specific expression after 36-40h APF (maintained at 27°C); Act88F-Gal4 (Bryantsev et al., 2012a) is a driver with IFM specific expression after 24h APF (maintained at 25°C) and Act79B-Gal4 (Dohn and Cripps, 2018) is a driver with TDT specific expression (maintained at 27°C). Temperature sensitive *Tubulin-Gal80^ts^*, as noted in figure panels and legends, was used to restrict some knockdown experiments to adult muscle development by a temperature shift of late third instar larvae from 18 °C to 29 °C. Rbfox1 over-expression with UH3-Gal4 was induced 40h APF onwards to avoid lethality at earlier stages.

### Behavioral assays

Flight behavior was tested as described previously (Drummond et al., 1991), or by introducing 30 adult males flies into a 1-meter long cylinder divided into 5 zones (Schnorrer et al., 2010). Flies landing in the top two zones are ‘normal fliers’, in the middle two zones are ‘weak fliers’ and at the bottom are ‘flightless’. Pupal eclosion (survival) was determined by counting the number of flies that eclose from at least 50 pupae of the appropriate genotype. Climbing ability was assayed using a modified rapid iterative negative geotaxis (RING) approach (Nichols et al., 2012). Adult males were collected on CO_2_ and recovered at least 24 hours before testing 3 times with a 1-minute recovery period for their ability to climb 5 centimeters in a 3 second or 5 second timeframe. Jumping ability was assayed as described previously (Chechenova et al., 2017). After clipping the wings and 24 hours recovery, 10-15 males were individually placed on A4 paper and gently pushed with a brush to stimulate the jump response. The start and the landing points were marked and the distance was calculated in centimeters.

### Rabbit anti-Bruno1 antibody generation

The divergent domain (DIV) region of Bru1 was cloned using SLIC into pCOOFY4 to generate His6-MBP-DIV. Primers are available in Supplemental Table 2. Fusion to MBP was necessary to maintain solubility. The protein was expressed in *E. coli* BL21-RIL cells and induced with 0.2 mM IPTG at 60 °C overnight. Expressed protein was purified over Ni-NTA beads and then cleaved with HRV3C-protease. MBP was depleted by incubation with Amylose beads. Protein was then dialyzed in buffer (200 mM NaCl, 50 mM Tris, 20 mM Imidazol) and sent as purified protein for antibody production (Pineda). Rabbit polyclonal antibodies were generated by Pineda according to a standard 120-day protocol. Resulting serum was affinity purified over an MBP column (to remove background antibodies generated against the MBP protein) followed by a column with beads coupled to Bru1-RA. Antibody bound to the column was eluted in citric acid and buffered to pH 7. Antibody was directly frozen in small aliquots in liquid nitrogen and stored at −80 °C until use.

### Immunofluorescence and microscopy

Fly hemi-thoraces were prepared for polarized microscopy as described previously (Nongthomba and Ramachandra, 1999). The hemi-thoraces were observed in an Olympus SZX12 microscope and photographed using Olympus C-5060 camera under polarized light optics. For confocal microscopy, flies were Bisected, fixed in 4% paraformaldehyde for 1h, washed with 0.3% PBTx (0.3% Triton X in PBS) for 15 min, and stained with 1:250 phalloidin-TRITC for 20 min. Sections were mounted on slides after washes with PBTx. Images were obtained using a Carl Ziess LSM 510 META confocal microscope.

Alternatively, IFMs and Abd-M were dissected and stained as previously described (Weitkunat and Schnorrer, 2014). All tissues were fixed for at least 30 minutes in 4% PFA in 0.5% PBS-T (1x PBS + Triton-X100). For visualization of IFMs, thoraces were cut longitudinally with a microtome blade. Abdominal muscle was fixed on a black silicon dissection dish, after the ventral part of the abdomen was carefully removed together with fat, gut and other non-muscle tissues. TDT (jump) muscle was exposed by opening the cuticle sagitally using fine biological forceps. One tip of the forceps was kept parallel to the fly thorax and gently inserted into the wing socket, allowing the initial split of the cuticle without damaging underlying tissues. The remaining cuticle covering the T2 mesothorax region, ventrally from the leg socket up to the dorsal bristles, was carefully removed to expose the underlying TDT muscle. Samples were blocked for 90 minutes at room temperature in 5% normal goat serum in PBS-T and stained with primary antibodies overnight at 4 °C. Rabbit anti-Bru1 1:500 and mouse anti-Lamin 1:100 (ADL67.10, DSHB) were used for staining. Samples were washed three times in 0.5% PBS-T for 10 min and incubated for 2 hours at room temperature with secondary conjugated antibodies (1:500) from Invitrogen (Molecular Probes), including Alexa488 goat anti-rabbit IgG, Alexa647 goat anti-mouse IgG, and rhodamine-phalloidin. Samples were washed three times in 0.5% PBS-T and mounted in Vectashield containing DAPI.

Confocal images were acquired on a Leica SP8X WLL upright using Leica LAS X software in the Core Facility Bioimaging at the Biomedical Center of the Ludwig-Maximilians-Universität München. Whole fly thorax images were taken with a HCPL FLUOTAR 10x/0.30 objective and detailed sarcomere structure was imaged with a HC PL APO 63x/1.4 OIL CS2 objective. Bru1 signal intensity was recorded at the same laser gain settings adjusted on the brightest control sample for each muscle type individually. All samples of same replicate were stained with the same antibody mix on the same day and imaged in the same imaging session.

### RNA isolation and RT-PCR

For *Rbfox1-*RNAi experiments, thirty flies were bisected and dehydrated in 70% ethyl alcohol overnight. IFM or TDT was dissected, homogenised and RNA isolated using TRI Reagent (Sigma) following the manufacturer’s instructions. RNA was confirmed using readings from Nanodrop software, and was converted to cDNA using a first strand cDNA synthesis kit (Fermentas, USA). Primers and PCR conditions are listed in Table S1.

For *Rbfox1*-IR^KK110518^ and *Rbfox1*-IR^27286^ experiments, IFM (from 30 flies) or TDT (from 60 flies) were dissected as previously described (Kao et al., 2019). For Abd-M, abdominal carcass was prepared from 15 flies in pre-cooled 1xPBS using fine biological forceps to remove fat, gut, trachea and other non-muscle tissues through a posterior cut in the abdomen. The abdomen was then removed from the thorax using fine scissors and snap-frozen in 50 μl of TRIzol (TRIzol Reagent, Ambion) on dry ice and immediately stored at −80^0^C. Dissection times were limited to maximum 30 minutes. RNA was isolated using the manufacturers protocol. Total RNA samples were treated with DNaseI (NEB) and measured on a Qubit 2.0 Fluorometer (Invitrogen). Comparable total RNA quantities were used for reverse transcription with LunaScript RT SuperMix Kit (New England Biolabs). cDNA was amplified with Phusion polymerase for 30-36 cycles and resulting PCR products were separated on a standard 1% agarose gel next to a 100 bp ladder (NEB). PCR primers are listed in Supplemental Table 2, with *Ribosomal protein L32* (*RpL32*, also called *RP49*) serving as an internal control in all reactions.

### RNA Immunoprecipitation (RIP) followed by cDNA synthesis

The RIP protocol was modified from (Carreira-Rosario et al., 2016). Approximately 500 mg of thoraces (from *Rbfox1^CC00511^* cultured flies) were lysed in 1 mL of RIPA buffer (50 mM Tris-HCl, 200 mM NaCl, 0.4% NP-40, 0.5% sodium deoxycholate, 0.1% SDS, 2 mM EDTA, 200 mM NaCl) with Sigma RNAse inhibitor, pre-cleared with Protein-G magnetic Dynabeads, and incubated with mouse anti-GFP (Developmental Studies Hybridoma Bank (DSHB), 12A6) or IgG isotype (purified from normal mouse serum). The beads with immunoprecipitated RNA bound to Rbfox1-GFP were washed and treated with Proteinase K (25 minutes in 37 °C), followed by a TRI-reagent based RNA extraction, cDNA synthesis and PCR as described above.

### Protein extraction and Western blotting

For *Rbfox1-*RNAi experiments, IFMs from 20 flies were dissected, “skinned”, and thin filaments extracted as previously described (Vikhorev et al., 2010). These samples were run on SDS-PAGE and transferred onto a nitrocellulose membrane (Milipore, product no. IPVH00010), using a semi-dry transfer apparatus. Blots were stained with rabbit anti-Actin or rabbit anti-TnI (1:1000; a gift from A. Ferrus) or mouse anti-Tubulin (1:1000, Sigma) and washed with TBS-Triton X (0.1%). Blots were incubated with HRP-conjugated secondary anti-rabbit or anti-mouse antibodies (1:5000 in TBS-Triton X), washed and developed on an X-ray film in the dark.

For *Rbfox1*-IR^KK110518^ and *Rbfox1*-IR^27286^ experiments, IFM from 8 flies, TDT from 20 flies or Abd from 6 flies was dissected as described above. Samples were homogenized in 20 μl of freshly made SDS-buffer (2% SDS, 240 mM Tris pH6.8, 0.005% Bromophenol blue, 40% glycerol, 5% β-mercaptoethanol), incubated at 95 °C for 3 min and stored at −20 °C. Samples were run on 10% SDS-PAGE for separation and then transferred onto nitrocellulose membranes (Amersham Protran 0.2 μm NC) for 2h at 120 V. Membranes were stained with Ponceau S (Sigma) to access the quality of the blotting. Membranes were de-stained and blocked with 5% non-fat milk solution in 0.5% Tween-TBS buffer (T-TBS) for 1h, washed and incubated for 1h at room temperature with primary antibodies (rabbit anti-Bru1, 1:500; rabbit anti-H2AZ, 1:2000). Membranes were washed three times with T-TBS for 15 min and incubated with goat anti-rabbit HRP-conjugated secondary antibodies (Bio-Rad) for 1 hour at room temperature. Following three rounds of washes, the membranes were developed using Immobilion Western chemiluminescent (Milipore) substrate and exposed to X-ray films (Fuji medical X-ray, Super RX-N).

### Co-immunoprecipitation and mass spectrometry

Approximately 500 mg of thoraces (from *Rbfox1^CC00511^* cultured flies) were lysed in 1 mL of RIPA buffer with Sigma protease inhibitor mix, pre-cleared with Protein-G magnetic Dynabeads (Thermo Scientific, 10030D), and incubated with mouse anti-GFP (DSHB, 12A6) or IgG isotype (purified from normal mouse serum). The beads with immunoprecipitated proteins bound to Rbfox1-GFP were washed in RIPA buffer, followed by protein elution and denaturation, as described previously (Carreira-Rosario et al., 2016). Proteins were analysed by SDS-PAGE and unique bands were cut and processed for mass spectrometric analysis following the protocol provided by the Proteomics facility, Molecular Biophysics Unit, Indian Institute of Science.

### Image analysis

Confocal image analysis was performed with Image J/Fiji (Schindelin et al., 2012). For every experiment 10 to 15 images were acquired from at least 10 individual flies. Fiber detachment was scored from Z-stacks of whole thorax images. Sarcomere length and width were measured using MyofibrilJ (Spletter et al., 2018), https://imagej.net/MyofibrilJ) based on rhodamine-phalloidin staining. Analysis of Bru1 intensity was performed manually in Fiji from at least three nuclei per image. Analysis of semi-quantitative RT-PCR gels and Western Blots was performed using the ‘gel analysis’ feature in Fiji. *RpL32* and H2AZ were used as internal normalization controls for RT-PCR and Western analysis, respectively. Data were saved into Microsoft Excel. Plotting and statistical analysis were performed in GraphPad Prism 9.

### Bioinformatics

Rbfox1 has been identified to bind (U)GCAUG motifs in both vertebrates and *Drosophila* (Nazario-Toole et al., 2018; Pedrotti et al., 2015). To identify possible Rbfox1 targets in muscle, we first identified all TGCATG motifs in the genome using PWMScan (https://ccg.epfl.ch/pwmtools/). The BED output was converted to a GRanges object in R, and sequence locations mapping to intron, 5’-UTR or 3’-UTR regions (based on Flybase dmel_r6.38 annotation files) were isolated. Gene identifiers were assigned based on genomic coordinates, and sequences were filtered to match gene orientation (ie to retain sequences present in the transcribed pre-mRNA). Lists of genes with Rbfox1 sites in introns, 5’-UTR or 3’-UTR regions were then subjected to enrichment analysis using PantherDB (Mi et al., 2021), GOrilla (Eden et al., 2009), or with custom gene sets (Spletter et al., 2018). Plots were generated in R using packages listed in Supplemental Table 3.

mRNA-Seq data used in this manuscript has been published previously (Spletter et al., 2015; Spletter et al., 2018) and is available from GEO under accession numbers GSE63707, GSE107247 and GSE143430. Data was mapped with STAR to ENSEMBL genome assembly BDGP6.22 (annotation dmel_r6.32 (FB2020_01)), indexed with SAMtools and features counted with featureCounts. Downstream analysis and visualization were performed in R using the packages listed in Supplemental Table 3. Differential expression was analyzed with DESeq2 and DEXSeq, which additionally generated normalized counts values. Read-tracks were visualized on the UCSC Genome Browser. Splice junction reads were exported from STAR, and junction use for hand-selected events was calculated as: (number of reads for select junction D^1^A^x^) / (total number of reads D^1^A^1^ + D^1^A^2^ … + D^1^A^n^) * 100, where D = donor and A = acceptor. In this way we could determine the percent of junction reads from a given donor that use acceptor “x”, or swap A and D to determine the percent of junction reads from a given acceptor coming from donor “x”.

## Data availability

Raw numbers used to generate plots are available in Supplementary Table 4. mRNA-Seq data are publicly available from GEO with accession numbers GSE63707, GSE107247 and GSE143430.

## Acknowledgements

We sincerely thank L. S. Shashidhara, R. Cripps, A. Ferrus, Sonal Jaishwal, Frank Schnorrer, the Bloomington Drosophila Stock Centre (BDSC), the Vienna Drosophila Resource Center (VDRC), and the Drosophila Stock Facility at NCBS, Bangalore, India, for providing flies. MLS is grateful to Andreas Ladurner for generous support. We thank John Sparrow and Shao-Yen Kao for inputs on manuscript preparation and critical comments. We thank Sandra Esser for excellent technical assistance. We acknowledge the Core Facility Bioimaging at the LMU Biomedical Center (Martinsried, DE) for confocal imaging support. We acknowledge the Indian Institute of Science (IISc), the Department of Science and Technology (DST) (DST FIST, 2008 – 2013 Ref. No. SR/FST/LSII-018/2007), the University Grant Commission (UGC-SAP to MRDG: Ref. No. F.3-47/2009 (SAP-II) and the Department of Biotechnology (DBT), Govt. of India, (DBT-IISC Partnership Program for Advanced Research in Biological Sciences & Bioengineering Sanction Order No: DBT/BF/PRIns/2011-12/IISc/28.9.2012), the Deutsche Forschungsgemeinschaft (MLS, 417912216), the Center for Integrated Protein Science Munich (CIPSM) at the Ludwig-Maximilians-University München (MLS), the Deutsche Gesellschaft für Muskelkranke e.V. (MLS), and the International Max Planck Research School (EN) for financial assistance.

## Author Contributions

Contributions are defined using CRediT role terminology (https://casrai.org/credit/).

Investigation (EN, KK, AM, CB),

Validation (EN, AM),

Writing – original draft (MLS, KK, UN),

Writing – review & editing (EN, MLS, KK, AM, CB, UN),

Conceptualization (MLS, UN),

Formal analysis (MLS, KK, EN, AM),

Visualization (MLS, EN, KK, AM),

Supervision (MLS, UN),

Funding acquisition (MLS, UN)

