## Supplemental Legends for "Rbfox1 is required for myofibril development and maintaining fiber-type specific isoform expression in *Drosophila* muscles"

**Supplemental Information**

**Supplemental Figure Legends**

**Supplemental Figure 1: Rbfox1 is differentially expressed between myofiber types and necessary for tubular muscle development.**

**A)** Scheme of *Rbfox1* gene region illustrating different isoforms (exons, red; UTR, black) and sequences targeted by the hairpins used in this manuscript (*Rbfox1-*RNAi, yellow; *Rbfox1-IR^27286^*, orange; *Rbfox1-IR^KK110518^*, magenta). Not drawn to scale. Normalized read counts from mRNA-Seq data show *Rbfox1* expression levels in IFM from *w^1118^* (blue), *bru1-IR* (light blue), and *salm* mutants (green), as well as tubular jump muscle (TDT, light green) and whole legs (dark green). **B)** Splice junction reads from mRNA-Seq data in A show preferential use of *Rbfox1* exons in fibrillar and tubular muscle. Data presented as percent of junction reads supporting a given splice event as diagrammed on the right for exons 7, 12, 14/15 and 17/18. **C**) Fold-change in *Rbfox1* expression between IFM and tubular muscle from semi-quantitative RT-PCR of *w^1118^*. Data normalized to *RpL32* expression levels. **D-E)** Knockdown efficiency in IFM with *Rbfox1-*RNAi (D) from RT-qPCR and *Rbfox1-IR^27286^* and *Rbfox1-IR^KK110518^* (E) from semi-quantitative RT-PCR. Significance in C, E determined by ANOVA and post-hoc Tukey and in D by paired t-test (not significant, ns; *= p < 0.05, **= p < 0.01, ***= p < 0.001), error bars indicate standard deviation.  **F-O)** Single-plane confocal images in the center of tubular TDT (F-J) and Abd-M (K-O) muscles showing myofibril (phalloidin stained actin, magenta) and nuclear (DAPI, green) arrangement in *w^1118^*, *bru1^M2^*, *Rbfox1-IR^27286^*, Dcr2 enhanced *Rbfox1-IR^27286^* and *Rbfox1-IR^KK110518^* genotypes. Myofibrils invade the space between nuclei after Rbfox1 knockdown (white arrows). **P-Q)** Examples of severe phenotypes with complete loss of myofibril structure in TDT (P) and Abd-M (Q) with *Rbfox1-IR^KK110518^*. Scale bars = 5 μm.

**Supplemental Figure 2: Rbfox1 is necessary for development of fibrillar IFM.**

**A-B)** Quantification of sarcomere length (A) and myofibril width (B) in IFM at 90 h APF and 1 day adult with *Rbfox1* knockdown and in *bru1^M2^* mutants. Significance determined by ANOVA and post-hoc Tukey (not significant, ns; ***= p < 0.001).  **C-D)** Single-plane confocal images showing normal sarcomere structure (phalloidin stained actin, greyscale) in both control (C) and Act88F-Gal4 driven *Rbfox1-IR^KK110518^* (D). Scale bars = 5 μm. **E)** Polarized light microscopy (E) of hemithoraxes with UH3-Gal4 driven overexpression of Rbfox1 reveals torn IFM myofibers (yellow arrowhead). A single-plane confocal micrograph (E’) showing thin and torn myofibrils (yellow arrows) with short sarcomeres in Rbfox1 OE IFM. **F-H)** Confocal projections of hemithoraces showing IFM myofiber structure in control (F) and *Rbfox1* knockdown using the deGradFP system (G, H). *Rbfox1CC00511- deGradFP* flies have torn (red arrow) and fewer intact myofibers (asterisks). **I-J)** Confocal images of *Rbfox1CC00511-deGradFP* flies (J) show actin accumulation (arrow in J) and loss of sarcomere structure (arrowhead in J) as compared to the control (I), and confirm deGradFP efficiency (GFP expression in I marked by arrows). Full genotypes: control, (pUASP1-deGradFP / CyO ; Rbfox1CC00511 / TM6, Tb) and knockdown, *Rbfox1CC00511-deGradFP*,  (pUASP1-deGradFP / CyO ; Rbfox1CC00511/ Mef2-Gal4). Scale bars = 2 μm.

**Supplemental Figure 3: Bioinformatic analysis of genes with Rbfox1 binding motifs and identification of putative target genes in muscle.**

**A)** Workflow and summary of bioinformatic search for the Rbfox1 canonical binding motif (TGCATG) in annotated intron (red), 5’-UTR (blue) and 3’-UTR (magenta) regions. **B)** Genes that contain at least one Rbfox1 binding motif classified on the basis of Molecular Function gene ontology (GO) terms in PANTHER-DB software. **C)** Percent of genes in a category with a Rbfox1 motif in an intron (red), UTR region (yellow) or both regions (orange). Categories include RNA-binding proteins (RBPs), transmembrane proteins (TMs), transcription factors (TFs), sarcomeric proteins (SPs), genes identified to have an RNAi phenotype in muscle (Schnorrer et al., 2010) (Muscle phenotype), and genes identified to be fibrillar muscle specific (Spletter et al., 2015) (Fibrillar genes). Category membership from (Spletter et al., 2018). N denotes the total number of genes in each category. **D)** Select biological process GO term enrichments in muscle phenotype genes with an Rbfox1 binding motif (full analysis available in Supplemental Table 1). **E-H)** Schematic of the *wupA* (E), *exd* (F), *Mef2* (G) and *bru1* (H) gene regions and annotated splice isoforms. Rbfox1 binding motifs are marked with vertical blue lines. Intronic binding site, magenta arrowhead; UTR binding site, green arrowhead; exon, red box; UTR, black box; RT-PCR primers, green arrows. Diagrams not drawn to scale.

**Supplemental Figure 4: Rbfox1 regulates mRNA levels of target genes and interacts with translation and NMD factors.**

**A-B)** RT-qPCR quantification of fold change in mRNA expression levels of *wupA* (A) and *Act88F* (B) in IFM and TDT from *Rbfox1*-RNAi and Rbfox1 OE. Normalized against *RpL32* signal. Significance is from paired t-test (not significant, ns; ** = p < 0.01). **C-D)** Semi-quantitative RT-PCR gel images (C) and quantification (D) for *Act88F* levels in IFM and TDT from *Rbfox1-IR^27286^* and *Rbfox1-IR^KK110518^*. Significance determined by ANOVA and post-hoc Tukey (not significant, ns; *= p < 0.05), error bars show standard deviation. **E)** SDS gel showing bands from input, pre-cleared lysate, proteins immunoprecipitated using IgG isotype antibody (control) and proteins immunoprecipitated using anti-GFP antibody from the thoraces of the *Rbfox1-GFP* (*Rbfox1CC00511*) line. Numbers 1 and 2 indicate immunoprecipitation from 2 biological repeats. Unique bands in the IP sample were cut and processed for mass spectrometric analysis. **F)** Peaks showing m/z ratios using MALDI-TOF. **G)** Possible interacting partners of Rbfox1 with a high Protein score include eIF4a and Rent1. *Protein score is -10*log(P), where P is the probability that the observed match is a random event. Scores > 50 are significant (p<0.05). **H)** Gel showing RT-PCR amplification of *bru1* (red arrowhead) from RIP using the *Rbfox1^CC00511^* line. **I)** Schematic of *bru1* (also called *arrest*) reporter-GFP construct. The promoter region containing 2.8 kilobases of sequence upstream of *bru1-RA* was cloned into the Ph-stinger vector (gel images). **J)** Representative Western blots and quantification of GFP reporter expression showing no significant change between control and *Rbfox1*-RNAi IFM.

**Supplemental Figure 5: Rbfox1-mediated regulation of Bru1 is expression level dependent.**

**A)** Diagram of the *Rbfox1*, *bru1*, and *mbl* gene loci (exons, red; UTR, black), which are all enriched for Rbfox1 binding motifs (blue lines). **B)** Standard normal count values for *Rbfox1* (magenta) and *bru1* (blue) from an mRNA-Seq developmental timecourse of wildtype IFM (Spletter et al., 2018). *Rbfox1* and *bru1* have opposite temporal expression profiles until 72 h APF. **C)** Quantification of RT-qPCR data for *bru1* transcript levels in IFM from *Rbfox1-*RNAi (left) or Rbfox1 OE (right). Significance is from paired t-test (** = p < 0.01; *** = p < 0.001). **D)** Quantification of fold change levels in *bru1* expression from semi-quantitative RT-PCR of IFM, TDT and Abd from control, *Rbfox1-IR^27286^* or Dcr2-enhanced *Rbfox1-IR^27286^* flies. PCR was performed with N-terminal (7 + 8) or C-terminal (14 + 17) primers common to all *bru1* isoforms or with *bru1-RB* specific primers (5 + 8), and normalized to *RpL32* expression level. **E-F)** Representative Western blot image (E) and quantification (F) for Bru1 in IFM, TDT and Abd showing a trend towards higher expression in *Rbfox1-IR^27286^* flies. H2AZ was used as a loading control and for intensity normalization. Significance in D, F determined by ANOVA and post-hoc Tukey (not significant, ns; * = p < 0.05, * = p < 0.01, *** = p < 0.001). **G)** Differential expression of *Rbfox1* in mRNA-Seq data based on DESeq2 comparison of IFM versus TDT (1 d adult), IFM versus *salm*^-/-^ IFM (1 d adult), IFM versus *bru1-IR* IFM (30 h APF, 72 h APF, 1 d adult), and IFM versus *bru1^M3^* IFM (not significant, ns; * = p < 0.05). **H)** RT-qPCR quantification of fold change in *Rbfox1* transcript level from IFM with Mef2-Gal4 driven Bru1 OE. Significance is from paired t-test (* = p < 0.05). **I-J)** Representative gel images (left) and quantification (right) of fold change in *Rbfox1* mRNA expression levels from semi-quantitative RT-PCR of IFM (I) and TDT (J) from *bru1^M2^* or Bru1 OE flies. Band intensity was normalized to *RpL32*. Significance determined by ANOVA and post-hoc Tukey (not significant, ns).

**Supplemental Figure 6: Rbfox1 and Bru1 genetically interact to regulate alternative splicing and muscle contractility.**

**A-D)** Confocal projections of hemithoraces showing phalloidin-stained IFMs of *Canton-S*, *Rbfox1*-RNAi, *bru1-IR*, and double *Rbfox1*-RNAi, *bru1-IR* knockdown. Scale bars = 100 μm. **E-P)** Single-plane confocal images of myofibril and sarcomere structure of the IFMs (E-H), TDT (I-L) and Abd-M (M-P) of *Canton-S*, *Rbfox1*-RNAi, *bru1-IR*, and double *Rbfox1*-RNAi, *bru1-IR* knockdown. Torn myofibers or myofibrils, arrowheads; abnormal actin accumulations, arrows. Scale bars = 10 μm. **Q-S)** Polarized microscopy images (Q-S) and single-plane confocal images (Q’-S’) showing IFM myofiber and myofibril structure in control (Q, Q’), in Mhc-Gal4 driven Bru1-PA overexpression (R, R’) or with Bru1 overexpression in an *Mhc^P401S^* mutant background (S, S’). Myofiber tearing (yellow arrowheads in R) and myofibril rupture (yellow arrows in R’) observed with Bru1 OE are partially rescued by using the Mhc^P401S^ allele. “Z” marks the Z-disc. Scale bars = 2 μm. **T)** Top: Diagram of the C-terminal region of the *wupA* locus illustrating the IFM isoform containing exon 4 and the tubular isoform that skips exon 4. One of exons 7, 8, 9 or 10 (dotted boxes) is alternatively spliced into *wupA* transcripts. Exons, yellow; UTR, tan, RT-PCR primers, black arrows. Bottom: Representative RT-PCR gel image with a primer specific to the *wupA-Ex4* isoform from IFM, TDT and Abd in genotypes as labelled*.* **U)** Representative RT-PCR gels for fiber-type specific alternative splice events in *Strn-Mlck* and *sls* in IFM, TDT and Abd, and an example of *RpL32* expression included as a control in all RT-PCR experiments. Diagrams of alternative exons and primer locations are on the right.

**Supplemental Figure 7: Rbfox1 regulates myogenic transcription factors including *Mef2* and *Salm*.**

**A)** Differential expression of *Mef2* and *Mef2* 5’-UTR exons 17, 20 and 21 in mRNA-Seq data from control IFM versus TDT (dark green), leg (light green), tubular-converted *salm^-/-^* IFM (green), *bru1^M3^* mutant IFM (blue) or *bru1-IR* IFM (light blue). log_2_(fold change) and significance p-values (grey values above bars) are from DESeq2 for overall *Mef2* transcript levels (pan *Mef2*) and from DEXSeq for relative exon use of *Mef2-Ex17*, *Mef2-Ex20* and *Mef2-Ex21*. Positive values show preference in wild-type IFM, while negative values show preference in tubular muscle or mutant IFM. **B)** Temporal and fiber-type selective mRNA-Seq splice junction use in the *Mef2* 5’-UTR region. Data presented as the percentage of junctions that use *Mef2-Ex17* (blue), *Mef2-Ex20* (magenta) or *Mef2-Ex21* (cyan) as the splice donor, and thus likely reflects the percent of transcripts that use the short 5’-UTR encoded by exon 17 versus the longer 5’-UTRs encoded by exons 20 and 21. Total junction reads for these events ranged from 22 to 1453, with an average of 334 ± 124 events per timepoint. **C)** Confocal images of thorax hemisections (C) and IFM myofibrils (C’) with overexpression of Mef2 driven by Mhc-Gal4. Myofibrils show actin accumulations (red arrow), but no hypercontraction. ‘*’ indicates IFM myofibers. **D)** Western blot demonstrating increased expression levels of Actin88F and TnI in IFM with Mef2 OE. **E)** RT-PCR confirmation of *Mef2* overexpression with Mhc-Gal4. Significance is from paired t-test (*** = p < 0.001). **F-G)** Single-plane confocal images from the surface (F, G) or center (F’, G’) of TDT from control (F) or *salm^-/-^* mutant (G) flies. Mild myofibril organization defects (yellow arrows) are observed in mutant TDT (phalloidin-stained actin, magenta; DAPI-stained nuclei, green). Scale bar = 5 μm. **H)** Confocal image of *salm-IR* IFM showing a tubular morphology. Scale bar = 2 μm. **I)** Confirmation of *salm* knockdown by semi-quantitative RT-PCR. **J)** RT-qPCR showing *bru1* levels are down regulated in *salm-IR* IFM. **K)** Plots showing quantification of the phenotypes shown in Fig. 7 H-P (N ≤ 28; 3 biological repeats).

**Supplemental Table 1: Bioinformatic identification of putative Rbfox1 targets in *Drosophila* muscle.**

This spreadsheet contains several tables of data for genes that contain TGCATG binding motifs in the *Drosophila* genome, including the raw output from PWMScan. We provide a table of the coordinates of all Rbfox1 binding sites in introns, 5’-UTR or 3’-UTR regions, as well as a list of genes and the number of Rbfox1 binding sites they contain. We also provide the full GO enrichment analysis from GOrilla and PantherDB.

**Supplemental Table 2: Primer Sequences**

This spreadsheet contains sequences of all primers used in this manuscript for PCR, RT-PCR, RIP and cloning purposes.

**Supplemental Table 3: Key Resources Table**

Here we include a table of all reagents used in this work, complete with their order numbers and sources. The table includes genes studied, fly lines, bacterial strains, plasmids, antibodies, chemicals and software.

**Supplemental Table 4: Raw Data**

This spreadsheet includes raw data used to generate plots throughout the manuscript.
