## Supplemental Figures for "Rbfox1 is required for myofibril development and maintaining fiber-type specific isoform expression in *Drosophila* muscles"

**Figure S1**

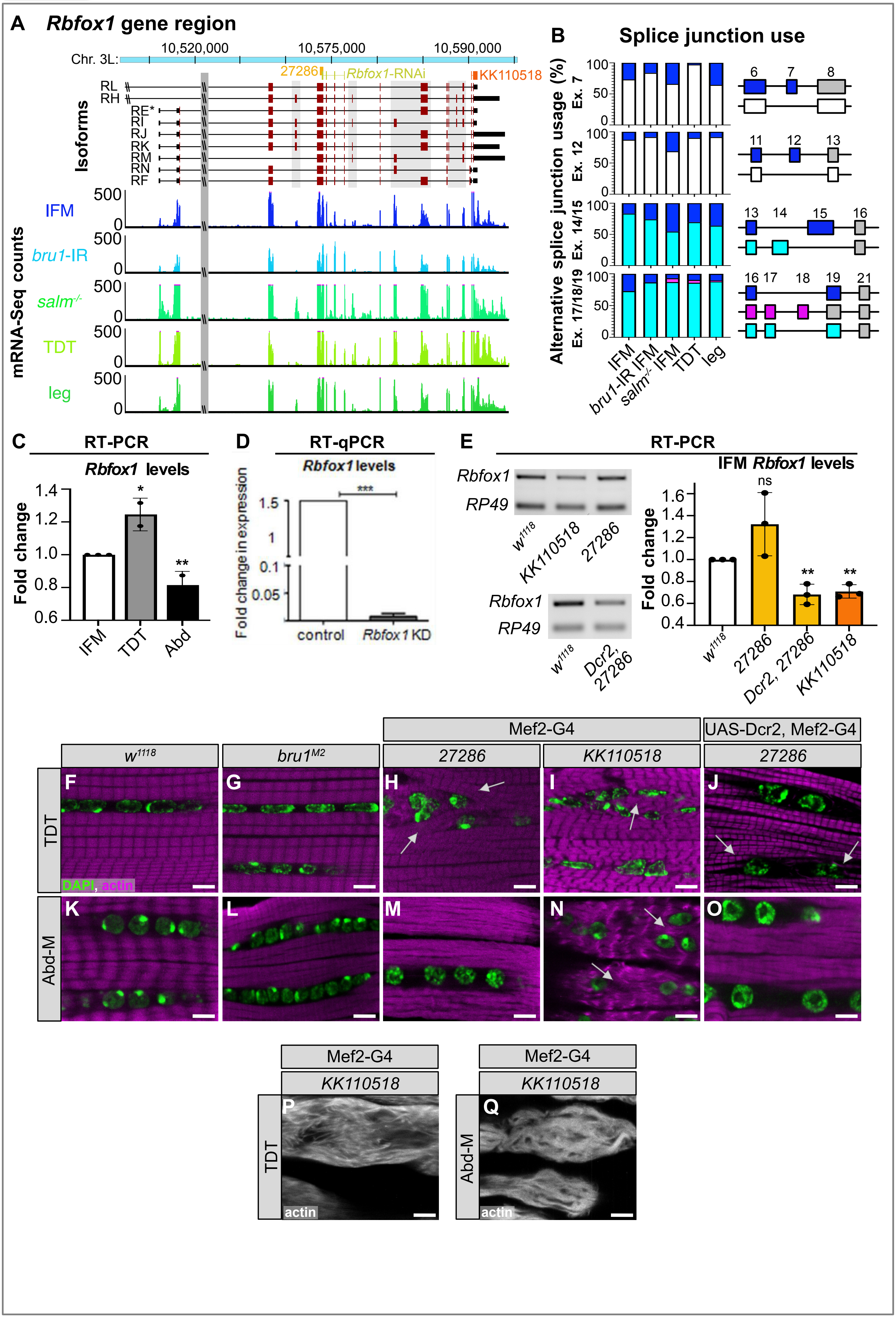

A Rbfox1 gene region
Chr. 3L:
10,520,000
10,575,000
10,590,000
27286
Rbfox1-RNAi
KK110518
Isoforms
RL
RH
RE*
RI
RJ
RK
RM
RN
RF
mRNA-Seq counts
IFM
bru1-IR
salm-/-
TDT
leg
500
0
B Splice junction use
Alternative splice junction usage (%)
Ex. 7
Ex. 12
Ex. 14/15
Ex. 17/18/19
100
50
0
IFM
bru1-IR IFM
salm-/- IFM
TDT
leg
6 7 8
11 12 13
13 14 15 16
16 17 18 19 21
C RT-PCR
Rbfox1 levels
Fold change
1.4
1.2
1.0
0.8
0.6
IFM
TDT
Abd
*
**
D RT-qPCR
Rbfox1 levels
Fold change in expression
1.5
1
0.1
0.05
***
control
Rbfox1 KD
E RT-PCR
Rbfox1
RP49
w1118
KK110518
27286
Dcr2, 27286
IFM Rbfox1 levels
Fold change
1.6
1.4
1.2
1.0
0.8
0.6
0.4
ns
**
27286
Dcr2, 27286
KK110518
Mef2-G4
UAS-Dcr2, Mef2-G4
w1118
bru1M2
TDT
Abd-M
F G H I J
K L M N O
DAPI, actin
Mef2-G4
KK110518
P
Q
actin

**Figure S2**

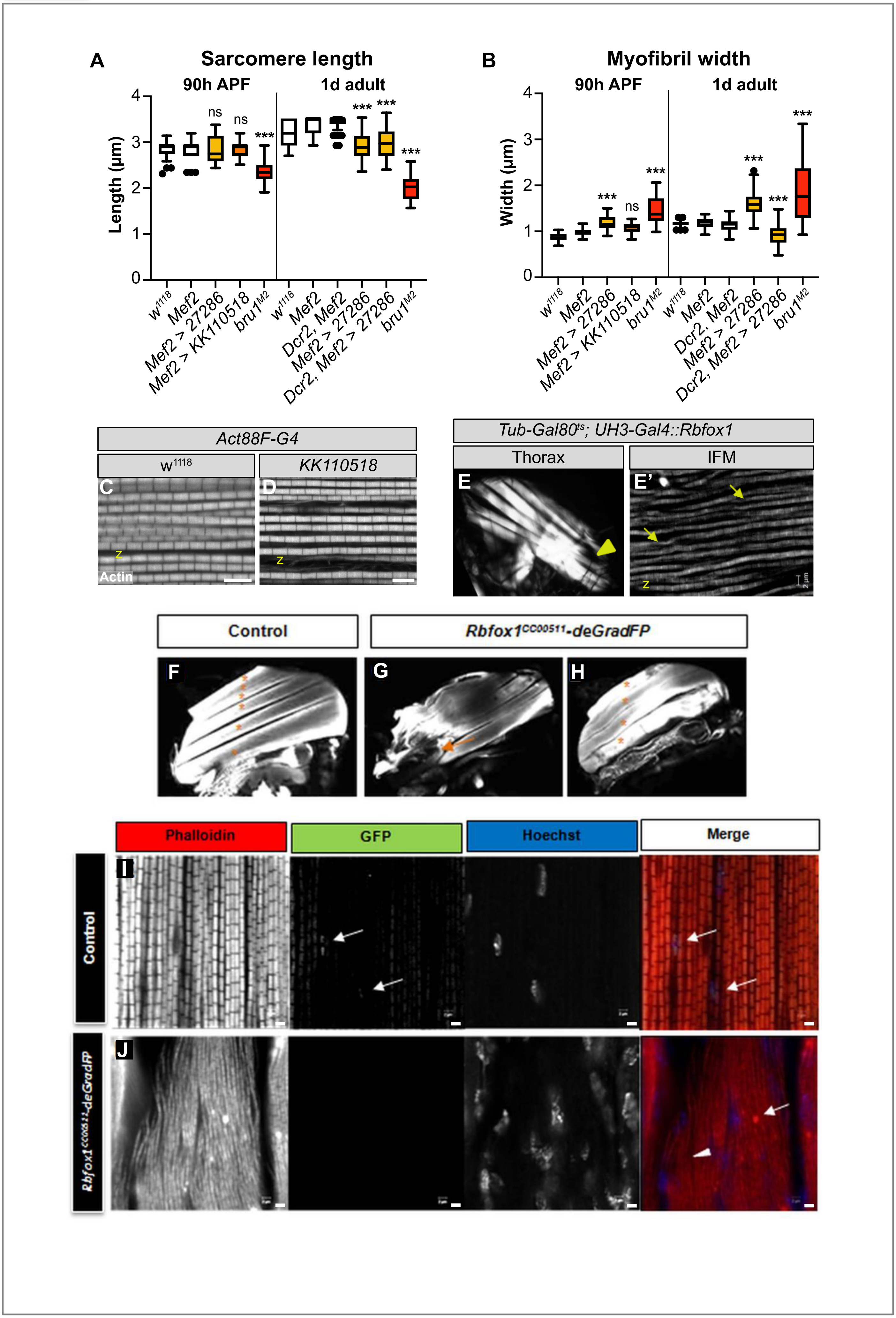

A
Sarcomere length
90h APF
1d adult
Length (µm)
ns
ns
***
w1118
Mef2
Mef2 > 27286
Mef2 > KK110518
bru1M2
Dcr2, Mef2
Dcr2, Mef2 > 27286
B
Myofibril width
Width (µm)
Act88F-G4
w1118
KK110518
C
D
Actin
z
Tub-Gal80ts; UH3-Gal4::Rbfox1
Thorax
IFM
E
E'
Control
Rbfox1CC00511-deGradFP
F
G
H
Phalloidin
GFP
Hoechst
Merge
I
J

**Figure S3**

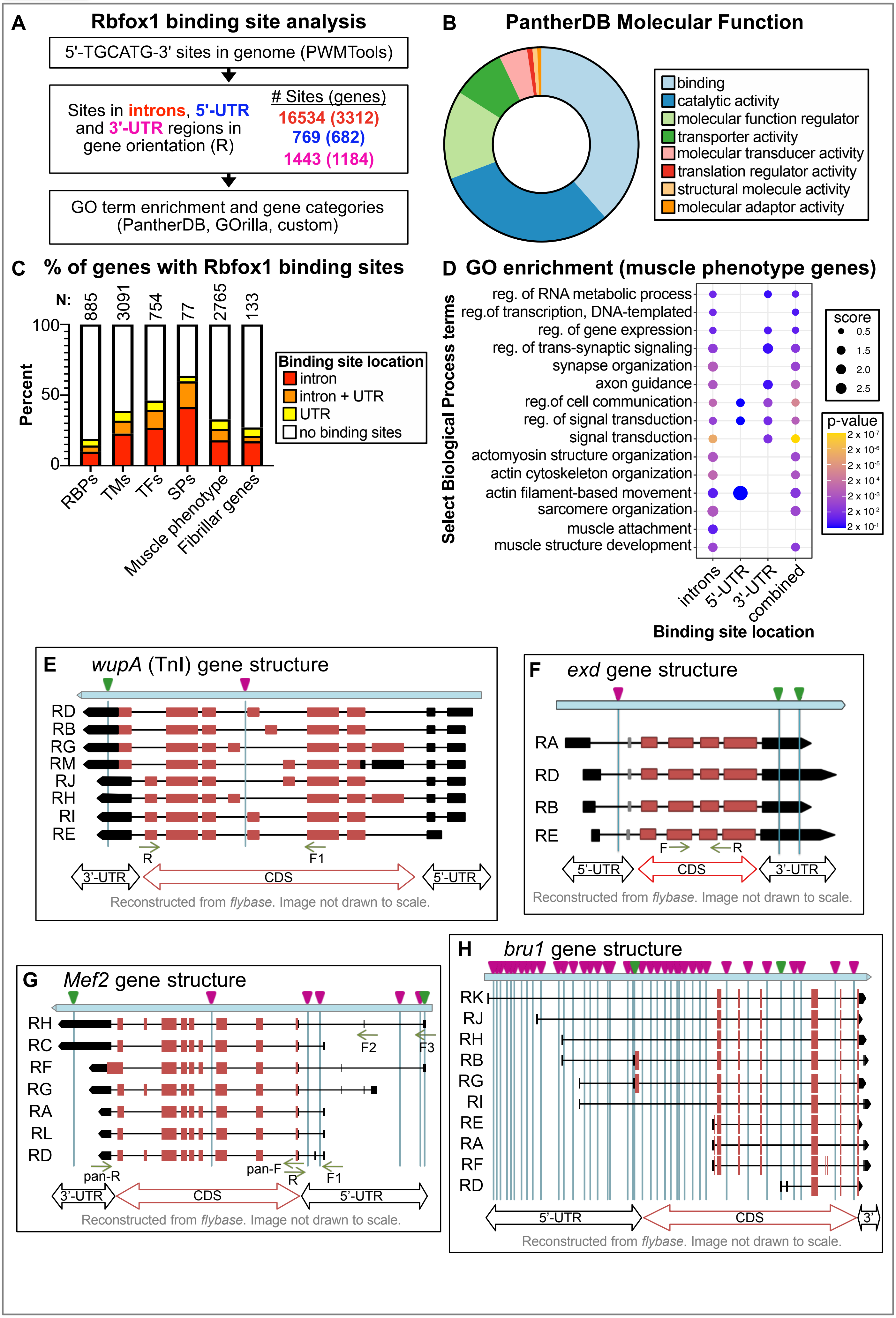
A
Rbfox1 binding site analysis
5'-TGCATG-3' sites in genome (PWMTools)
Sites in introns, 5'-UTR and 3'-UTR regions in gene orientation (R)
### Sites (genes)
16534 (3312)
769 (682)
1443 (1184)
GO term enrichment and gene categories (PantherDB, GOrilla, custom)
B
PantherDB Molecular Function
binding
catalytic activity
molecular function regulator
transporter activity
molecular transducer activity
translation regulator activity
structural molecule activity
molecular adaptor activity
C
% of genes with Rbfox1 binding sites
N: 885 3091 754 77 2765 133
Percent
100
50
0
RBPs
TMs
TFs
SPs
Muscle phenotype
Fibrillar genes
Binding site location
intron
intron + UTR
UTR
no binding sites
D
GO enrichment (muscle phenotype genes)
Select Biological Process terms
reg. of RNA metabolic process
reg.of transcription, DNA-templated
reg. of gene expression
reg. of trans-synaptic signaling
synapse organization
axon guidance
reg.of cell communication
reg. of signal transduction
signal transduction
actomyosin structure organization
actin cytoskeleton organization
actin filament-based movement
sarcomere organization
muscle attachment
muscle structure development
introns
5'-UTR
3'-UTR
combined
Binding site location
score
0.5
1.5
2.0
2.5
p-value
2 x 10-7
2 x 10-6
2 x 10-5
2 x 10-4
2 x 10-3
2 x 10-2
2 x 10-1
E
wupA (TnI) gene structure
RD
RB
RG
RM
RJ
RH
RI
RE
R
F1
3'-UTR
CDS
5'-UTR
Reconstructed from flybase. Image not drawn to scale.
F
exd gene structure
RA
RD
RB
RE
F
R
5'-UTR
CDS
3'-UTR
Reconstructed from flybase. Image not drawn to scale.
G
Mef2 gene structure
RH
RC
RF
RG
RA
RL
RD
F2
F3
pan-R
pan-F
R
F1
3'-UTR
CDS
5'-UTR
Reconstructed from flybase. Image not drawn to scale.
H
bru1 gene structure
RK
RJ
RH
RB
RG
RI
RE
RA
RF
RD
5'-UTR
CDS
3'
Reconstructed from flybase. Image not drawn to scale.

**Figure S4**

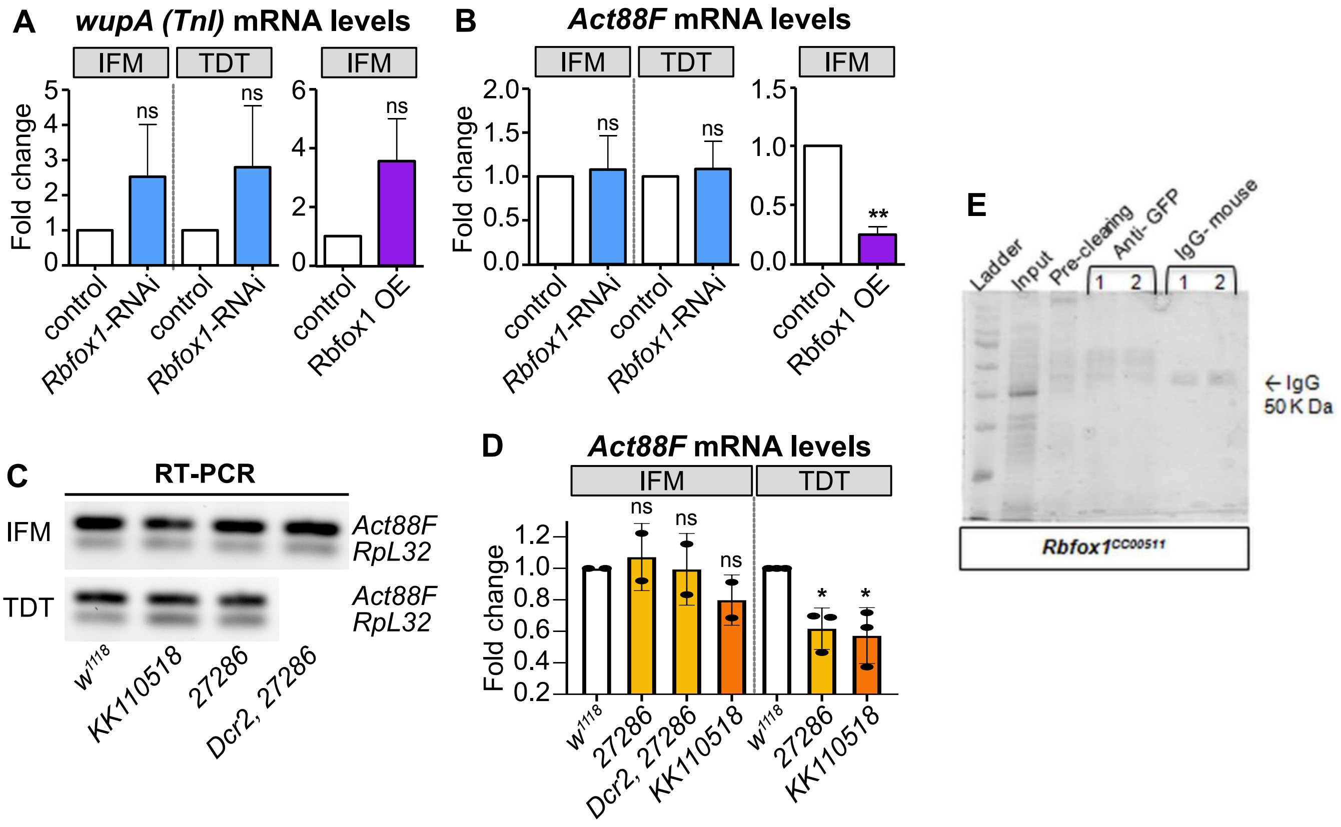


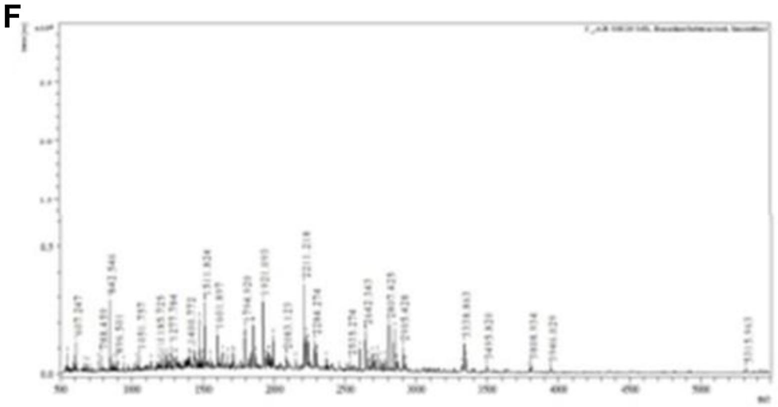


**G Interacting partners of Rbfox1**

| Name | Initiation factor (eIF4a) | Rent1 - Regulator of nonsense transcripts homolog 1 |
|---|---|---|
| Biological process | Translation initiation | Non-sense mediated decay |
| Mass | 13125 | 46249 |
| Protein Score* | 57 | 61 |
| Matches in biological repeats | 12 | 14 |

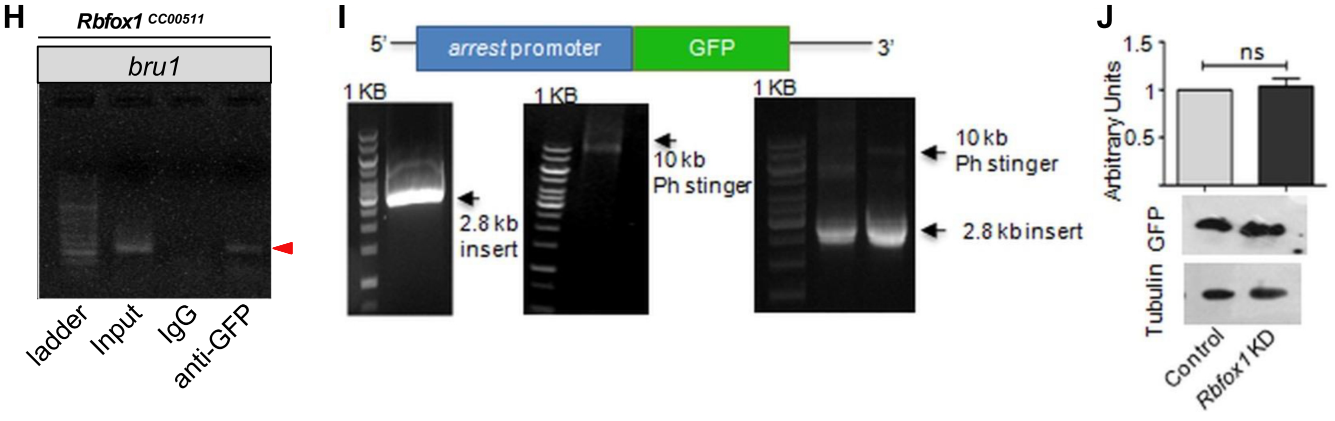

**Figure S5**

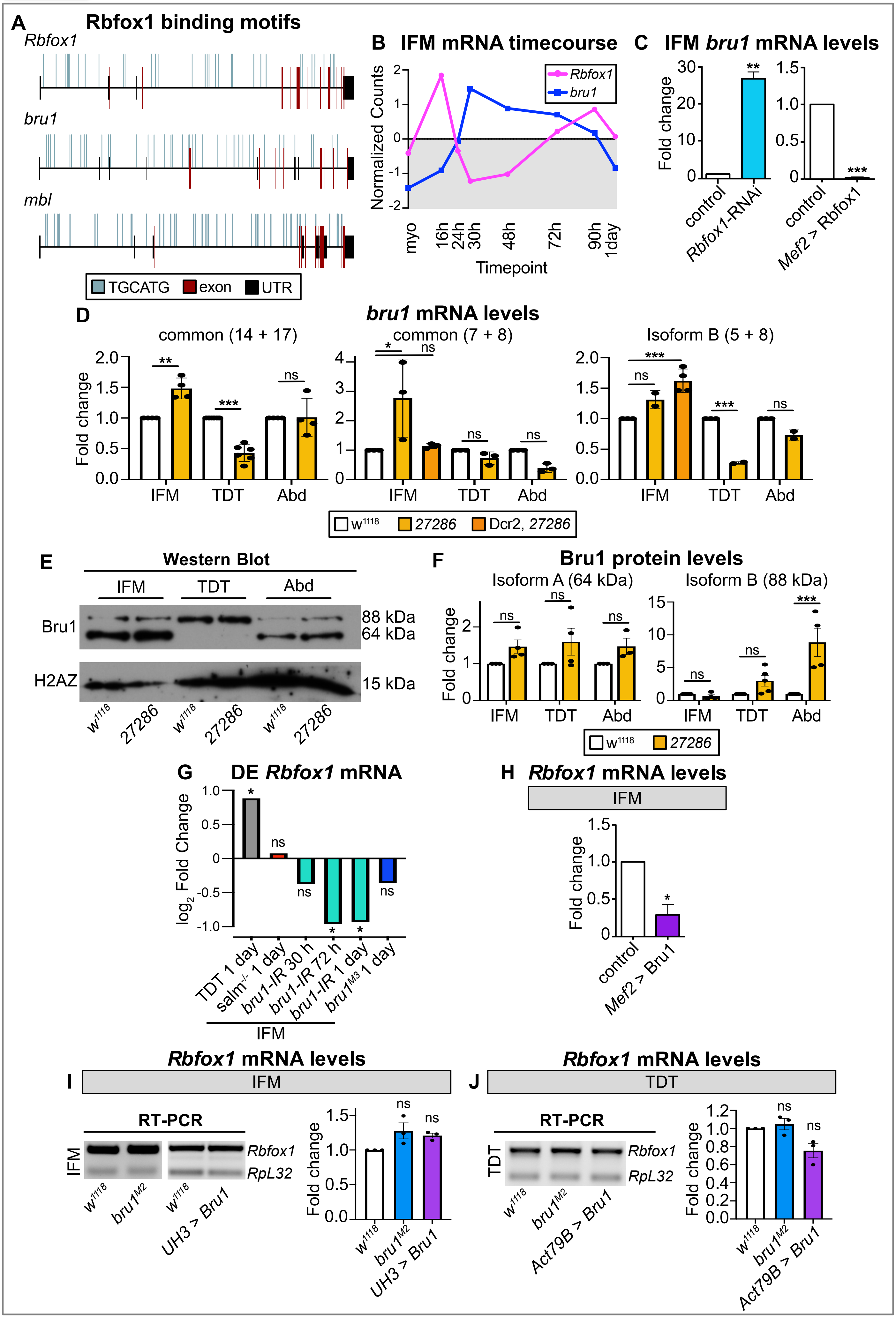
A
Rbfox1 binding motifs
Rbfox1
bru1
mbl
TGCATG
exon
UTR
B
IFM mRNA timecourse
Rbfox1
bru1
Normalized Counts
2
1
0
-1
-2
myo
16h
24h
30h
48h
72h
90h
1day
Timepoint
C
IFM bru1 mRNA levels
Fold change
30
20
10
0
**
control
Rbfox1-RNAi
1.5
1.0
0.5
0
***
control
Mef2 > Rbfox1
D
bru1 mRNA levels
common (14 + 17)
common (7 + 8)
Isoform B (5 + 8)
Fold change
2.0
1.5
1.0
0.5
0.0
4
3
2
1
0
**
***
*
***
ns
IFM
TDT
Abd
w1118
27286
Dcr2, 27286
E
Western Blot
IFM
TDT
Abd
Bru1
H2AZ
88 kDa
64 kDa
15 kDa
w1118
27286
F
Bru1 protein levels
Isoform A (64 kDa)
Isoform B (88 kDa)
Fold change
2
1
0
15
10
5
0
ns
***
IFM
TDT
Abd
w1118
27286
G
DE Rbfox1 mRNA
log2 Fold Change
1.0
0.5
0
-0.5
-1.0
*
ns
TDT 1 day
salm-/- 1 day
bru1-IR 30 h
bru1-IR 72 h
bru1-IR 1 day
bru1M3 1 day
IFM
H
Rbfox1 mRNA levels
IFM
Fold change
1.5
1.0
0.5
0
*
control
Mef2 > Bru1
I
Rbfox1 mRNA levels
IFM
RT-PCR
Rbfox1
RpL32
w1118
bru1M2
UH3 > Bru1
Fold change
1.5
1.0
0.5
0.0
ns
J
Rbfox1 mRNA levels
TDT
RT-PCR
Rbfox1
RpL32
w1118
bru1M2
Act79B > Bru1
Fold change
1.2
1.0
0.8
0.6
0.4
0.2
0.0
ns

Figure S6

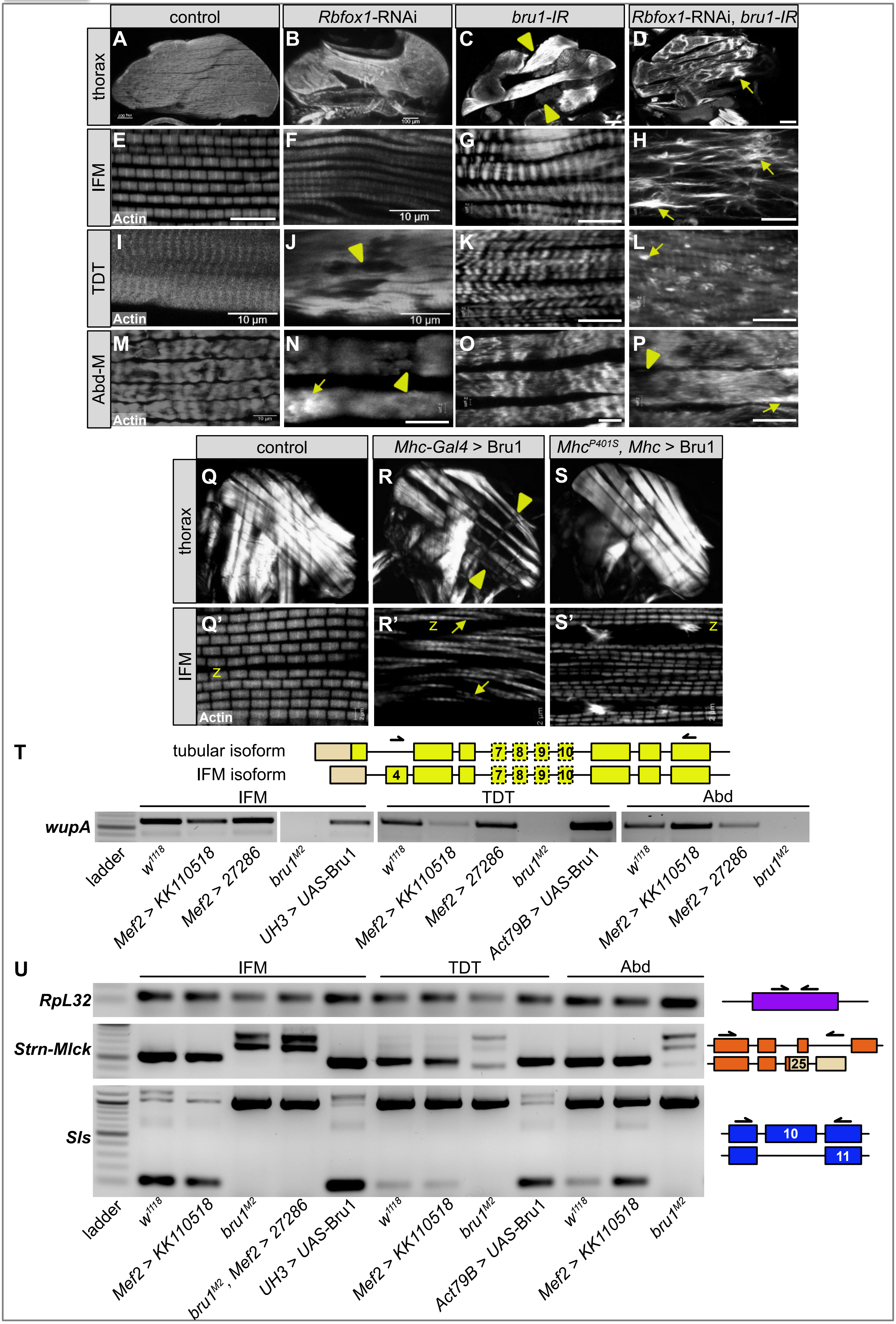

control
Rbfox1-RNAi
bru1-IR
Rbfox1-RNAi, bru1-IR
thorax
IFM
TDT
Abd-M
A
B
C
D
E
F
G
H
I
J
K
L
M
N
O
P
Actin
10 µm
100 µm
control
Mhc-Gal4 > Bru1
MhcP401S, Mhc > Bru1
Q
R
S
Q'
R'
S'
Z
T
tubular isoform
IFM isoform
4
7
8
9
10
IFM
TDT
Abd
wupA
ladder
w1118
Mef2 > KK110518
Mef2 > 27286
bru1M2
UH3 > UAS-Bru1
Act79B > UAS-Bru1
U
RpL32
Strn-Mlck
Sls
25
11
bru1M2, Mef2 > 27286

**Figure S7**

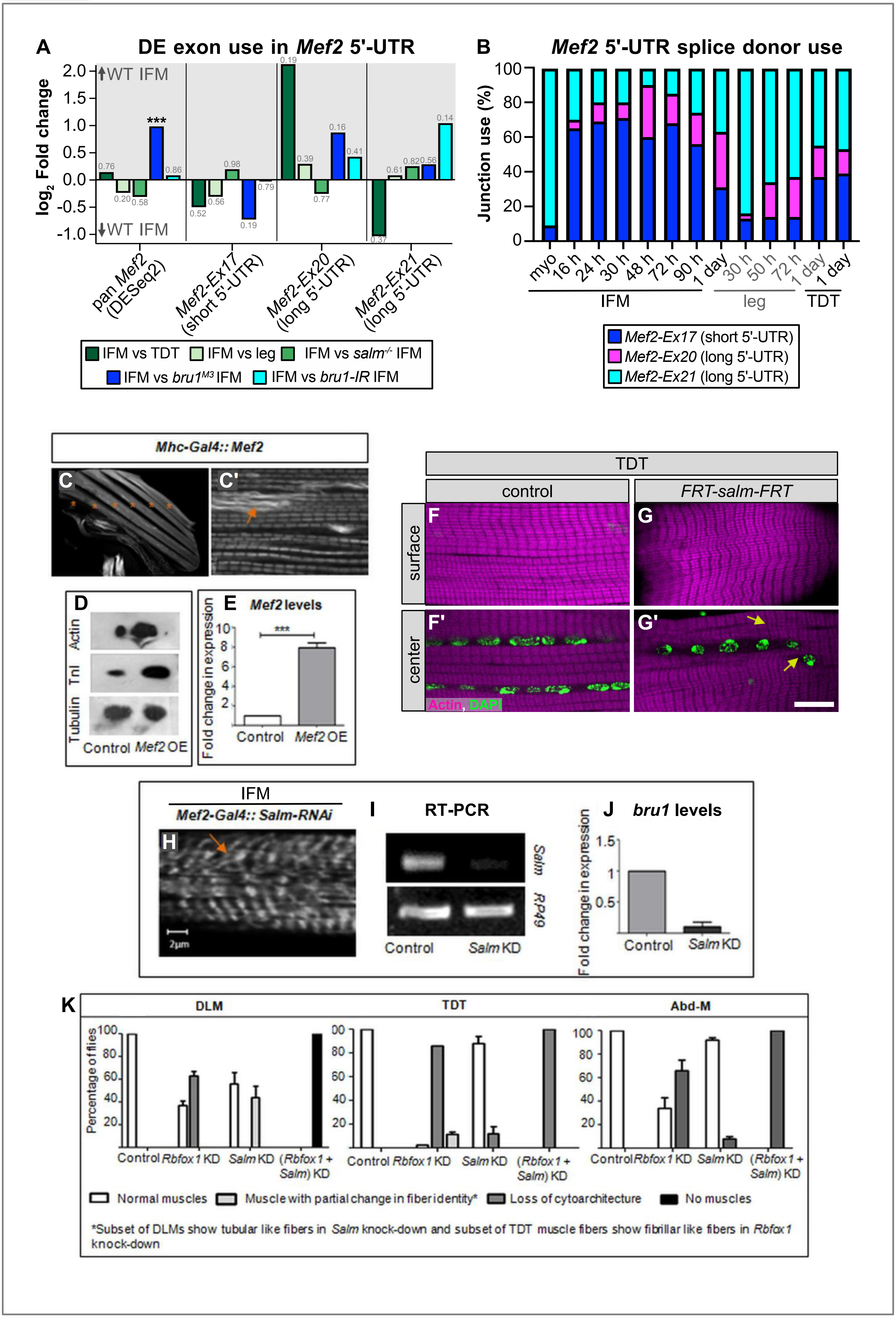
A
DE exon use in Mef2 5'-UTR
log2 Fold change
WT IFM
WT IFM
pan Mef2 (DESeq2)
Mef2-Ex17 (short 5'-UTR)
Mef2-Ex20 (long 5'-UTR)
Mef2-Ex21 (long 5'-UTR)
IFM vs TDT
IFM vs leg
IFM vs salm-/- IFM
IFM vs bru1M3 IFM
IFM vs bru1-IR IFM
B
Mef2 5'-UTR splice donor use
Junction use (%)
myo
16 h
24 h
30 h
48 h
72 h
90 h
1 day
30 h
50 h
72 h
1 day
1 day
IFM
leg
TDT
Mef2-Ex17 (short 5'-UTR)
Mef2-Ex20 (long 5'-UTR)
Mef2-Ex21 (long 5'-UTR)
Mhc-Gal4:: Mef2
C
C'
D
Actin
TnI
Tubulin
Control
Mef2 OE
E
Mef2 levels
Fold change in expression
Control
Mef2 OE
TDT
control
FRT-salm-FRT
surface
center
F
G
F'
G'
Actin, DAPI
IFM
Mef2-Gal4:: Salm-RNAi
H
2µm
I
RT-PCR
Salm
RP49
Control
Salm KD
J
bru1 levels
Fold change in expression
Control
Salm KD
K
DLM
TDT
Abd-M
Percentage of flies
Control
Rbfox1 KD
Salm KD
(Rbfox1 + Salm) KD
Normal muscles
Muscle with partial change in fiber identity*
Loss of cytoarchitecture
No muscles
*Subset of DLMs show tubular like fibers in Salm knock-down and subset of TDT muscle fibers show fibrillar like fibers in Rbfox1 knock-down
